# Parallel evolution and cryptic diversification in the common and widespread Amazonian tree, *Protium subserratum*

**DOI:** 10.1101/2022.09.28.509925

**Authors:** Tracy M. Misiewicz, Tracey Simmons, Benjamin E. Carter, Paul V. A. Fine, Abigail J. Moore

## Abstract

The lowland Amazon rainforest houses some of the greatest tree diversity on earth. While the vast majority of these species are rare, a small number are common and widespread and thus considered to play a disproportionate role in many of the global ecosystem services provided by the Amazon. However, the extent to which dominant Amazonian tree species actually include multiple clades, each on their own unique evolutionary trajectory, is unknown. Here we investigate the extent to which lineage divergence may be occurring within *Protium subserratum* (Burseraceae), a common and widespread tree species that is monophyletic with populations exhibiting genotypic and phenotypic differences associated with soil and geography. Utilizing a combination of phylogenomic and population genomic methods with sampling from across the range, we found that *P. subserratum* contains at least eight distinct clades. Specialization onto white-sand soils has evolved independently at least two times within the species; however, phenotype is not correlated with soil type. Finally, cryptic diversity at the base of the Andes is associated with elevational shifts. Together these results lend support to the hypothesis that common and widespread Amazon tree species may not represent evolutionary cohesive units. Instead, these dominant species may more commonly represent species complexes, undergoing evolutionary transitions on a trajectory to become multiple range restricted, specialist species.

## Introduction

Home to an estimated 16,000 tree species, the lowland Amazon rainforest houses some of the greatest diversity on earth (ter Steege et al., 2013). While the vast majority of these species are rare, a small number are common and widespread (ter Steege et al., 2013). For instance, ter Steege et al. (2013) estimated that only 227 tree species represent more than 50% of individual Amazonian trees with a dbh of >10cm. If true, these results suggest that just a tiny proportion of Amazonian tree taxa are disproportionally responsible for ecosystem process and biogeochemical cycles with global impacts. In turn, it has been suggested that understanding the ecology, biology and distribution of these common taxa should be prioritized given their outsized importance and their potential to serve as a proxy for understanding general patterns of biodiversity (ter Steege et al. 2013). Still, caution is warranted because evolutionary diversity housed within widespread and common taxa may go unrecognized for a number of reasons. Morphological and/or ecological variation may be undiscovered or lacking clear breaks, making delineation difficult. Additionally, recent divergence and/or introgression may lead to shared alleles or incomplete lineage sorting among clades, and taxa with large geographic ranges may be under sampled (Bickford et al., 2007). In Amazonian tree taxa these challenges are compounded by the fact that basic taxonomic, ecological and distributional data are often lacking (Cardoso et al., 2017; Crane, 2004; Goodwin et al., 2015) and sampling within species for molecular studies is sparse (Pennington & Lavin, 2016). As a result, our understanding of the extent to which evolutionary complexity may exist within what are considered to be single common and widespread tree species remains largely unknown.

To date, few studies have attempted to assess genetic structure within widespread Amazonian tree taxa. However, results across those that have are variable. Early studies suggested that two widespread Amazonian tree species do in fact represent cohesive evolutionarily taxonomic units, exhibiting weak genetic structure even over large geographic areas. For example, *Ceiba pentandra* (L.) Gaertn. (Malvaceae), the 744^th^ most abundant Amazonian tree species (ter Steege et al., 2013) exhibited three internal transcribed spacer (ITS) haplotypes from populations sampled from across the Amazon (Dick et al., 2007). Likewise, the 23^rd^ most abundant species, *Symphonia globulifera* L.f. (Clusiaceae) shared just one ITS haplotype across populations from French Guiana, Ecuador and Bolivia (Dick et al., 2003) and even when sampling was expanded to include populations more than 2500km apart F_ST_ values and ITS variation remained low (Dick & Heuertz, 2008).

Conversely, more recent studies have presented evidence that while some widespread tree species do represent monophyletic groups, they are not necessarily single evolutionarily cohesive units maintained by gene flow. Instead they are composed of multiple lineages each on their own independent evolutionary trajectory. For instance, the 22^nd^ most common Amazonian tree (ter Steege et al., 2013), *Carapa guianensis* Aubl. (Meliaceae), was found to include six distinct haplogroups that did not exhibit geographic structuring, suggesting the possible existence of cryptic lineages (Scotti-Saintagne et al., 2013). *Astrocaryum murumuru* Mart. (Arecaceae), historically defined as a single widespread species with geographically discrete varieties (Henderson et al., 1995), was ranked 10^th^ in the list of hyperdominant Amazonian trees (ter Steege et al., 2013). However, more recent molecular studies have redefined the species as *Astrocaryum* sect. *Huicungo* F. Kahn, a clade composed of 15 distinct species (Roncal et al., 2013, 2015). Similarly, *Protium heptaphyllum* Marchand (Burseraceae), ranked as the 12^th^ most abundant Amazonian tree (ter Steege et al., 2013), was recently proposed to include eight distinct taxa based on molecular, ecological and morphological data (Damasco et al., 2021). Given the variable nature of these results, more studies that include greater sampling within species are necessary to determine the degree to which widespread and common Amazonian trees actually include multiple evolutionary lineages.

For clades restricted to the lowland Amazon rainforest, diversification has been attributed to vicariance, dispersal, and adaptive divergence across habitat boundaries (Reviewed in Dick & Pennington, 2019). Major landscape level changes associated with the evolution of the modern-day Amazon river system are thought to have catalyzed speciation in the region by fragmenting previously continuous distributions of widespread species and hindering gene flow among populations (Nazareno et al., 2017; Smith et al., 2014; Wallace, 1854). Long distance dispersal events are also recognized for their importance in the establishment of geographically isolated populations (Dexter et al., 2017; Pennington & Lavin, 2016). Finally, habitat heterogeneity can facilitate lineage diversification, by promoting either divergence in allopatry if habitat is patchily distributed across the landscape or divergence in parapatry if selection across environmental gradients is strong (Nosil, 2012; Rundle & Nosil, 2005). The role of soils as a driver of diversification in Amazonian trees been the focus of numerous studies (Fine et al., 2014; Fine & Baraloto, 2016; Fine et al., 2005; Misiewicz et al., 2020; Misiewicz & Fine, 2014; Vicentini, 2016) and is considered to be an important factor in driving and maintaining tree diversity in the Amazon rainforest.

Amazonian soils vary widely in particle size, nutrient levels and their ability to retain moisture and thus are likely to exert strong selective pressures on the organisms that inhabit them (Pregitzer et al., 2010; Smith et al., 2012). White-sand soils are particularly distinct from other soils in the lowland Amazon. They are composed primarily of extremely nutrient poor white quartz sand and are patchily distributed across the Amazon (Adeney et al. 2016). White-sand forests range in area from just a few hectares to thousands of square kilometers and are notable for their high levels of tree endemism (Adeney et al., 2016; Fine et al., 2010; Guevara et al., 2016; Stropp et al., 2011; Vormisto et al., 2000). White-sand soils appear to have impacted lineage diversification of tropical trees in very different ways. For instance, the white-sand endemic genus *Pagamea* Aubl. (Rubiaceae), a widespread group with its diversity centered in the Guiana Shield, appears to have diversified via allopatric speciation driven by migration and subsequent geographic isolation across the patchy distribution of white-sand habitats (Vicentini, 2016). In the western Amazon, where white-sand soil patches are small (often <1 km^2^) and interspersed among comparatively young and more nutrient rich brown-sand and clay soils (Fine et al., 2005; Frasier, 2008; Hoorn, 1993), species within the genus *Protium* Burm. f. (Burseraceae) have evolved specialization onto white-sand soils multiple times suggesting that natural selection could play an important role in driving diversification within the group (Fine et al., 2014; Fine et al., 2005).

*Protium subserratum* (Engl.) Engl. is common and widespread across the Amazon Basin, ranked 264^th^ on the list of most abundant species by ter Steeg et al. (2013). The species represents a monophyletic lineage (Daly & Fine, 2011; Fine et al., 2014; Fine et al., 2013) and is notable both within the genus and the Amazonian tree flora, as one of the few species that both occurs in French Guiana and the western Amazon (Baraloto et al., 2021) and occupies multiple soil habitats with populations exhibiting genotypic and phenotypic differences associated with both soil and geography (Daly & Fine, 2011; Fine et al., 2014; Fine et al., 2005; Lokvam et al., 2015; Lokvam & Fine, 2012; Misiewicz & Fine, 2014). While the ecology and evolution of *P. subserratum* have been the focus of numerous studies, no single study has examined diversification within the species with adequate sampling across the genome and the range. Here we applied a combination of phylogenomic and population genomic methods with sampling from across the Amazon using herbarium specimens to address the following questions: 1) Does *P. subserratum* contain multiple independently evolving lineages? 2) How many times has specialization onto white-sand soils arisen? 3) Is phenotype correlated with soil type? and 4) Are the lineages within *P. subserratum* defined by ecology, geography or both?

### Study System

*Protium subserratum* is placed within section *Papilloprotium* Daly & P. Fine and is sister to *P. alvarezianum* Daly & P. Fine and *P. reticulatum* (Engl.) Engl., both white-sand specialists distributed across white-sand forest patches of the Rio Negro Basin of Venezuela and Brazil (Daly and Fine 2011). It has small, nectar producing, fragrant, white flowers and red fruits that are approximately 1.5 cm in diameter (Misiewicz, personal observation). Like most Amazonian trees, *P. subserratum* is dioecious. Stingless bees are the most likely pollinators (Misiewicz et al., 2020) and monkeys and large birds are hypothesized to be potential seed dispersers (Daly, 1987).

Both phenotypic and genotypic differences have been observed among geographically and ecologically disparate populations (Daly & Fine, 2011; Misiewicz & Fine, 2014). Daly and Fine (2011) utilized leaf and reproductive morphology to group *P. subserratum* individuals into four morphotypes, two morphotypes revolving around soil affiliation (non-white-sand and white-sand soils, morphotypes 2 and 3) and two corresponding with restricted geographic distributions (non-white-sand forests of French Guiana and eastern Brazil and non-white-sand forests restricted to Colombia’s Caquetá department, morphotypes 1 and 4). Phylogeographic analysis by Fine et al. (2013) utilized three nuclear genes and included individuals representing morphotypes 1, 2, and 3. While their results were suggestive of genetic structure within *P. subserratum* no groupings were resolved with support. Further population genetic analysis using microsatellites focused on populations found in the Department of Loreto, Peru and found strong evidence of genetic structuring around soil type (Misiewicz & Fine, 2014). Genetic groups corresponded to white-sand, brown-sand and clay soil types and little evidence of successful gene flow across edaphic boundaries was observed.

## Materials and Methods

### Taxon sampling

We sampled leaf material from 207 herbarium specimens collected between 1933 and 2018 from the New York Botanic Garden (NY), Missouri Botanic Garden (MO), the National Herbarium (US), and the University Herbarium at the University of California, Berkeley (UC) (Table S1). A total of 187 individuals of *Protium subserratum* were sampled from across the Amazon (Fig 1). For our outgroups, we sampled individuals from the remaining three species that, along with *P. subserratum,* comprise the section *Papilloprotium*. These included *P. alvarezianum* (5 individuals), *P. reticulatum* (5 individuals), *P. ferrugineum* (Engl.) Engl. (9 individuals) and *P. amazonicum* (Cuatrec.) Daly (1 individual). Usable sequences for outgroup species were only acquired for seven *P. ferrugineum* individuals and the single *P. amazonicum* individual.

**Figure 1:**
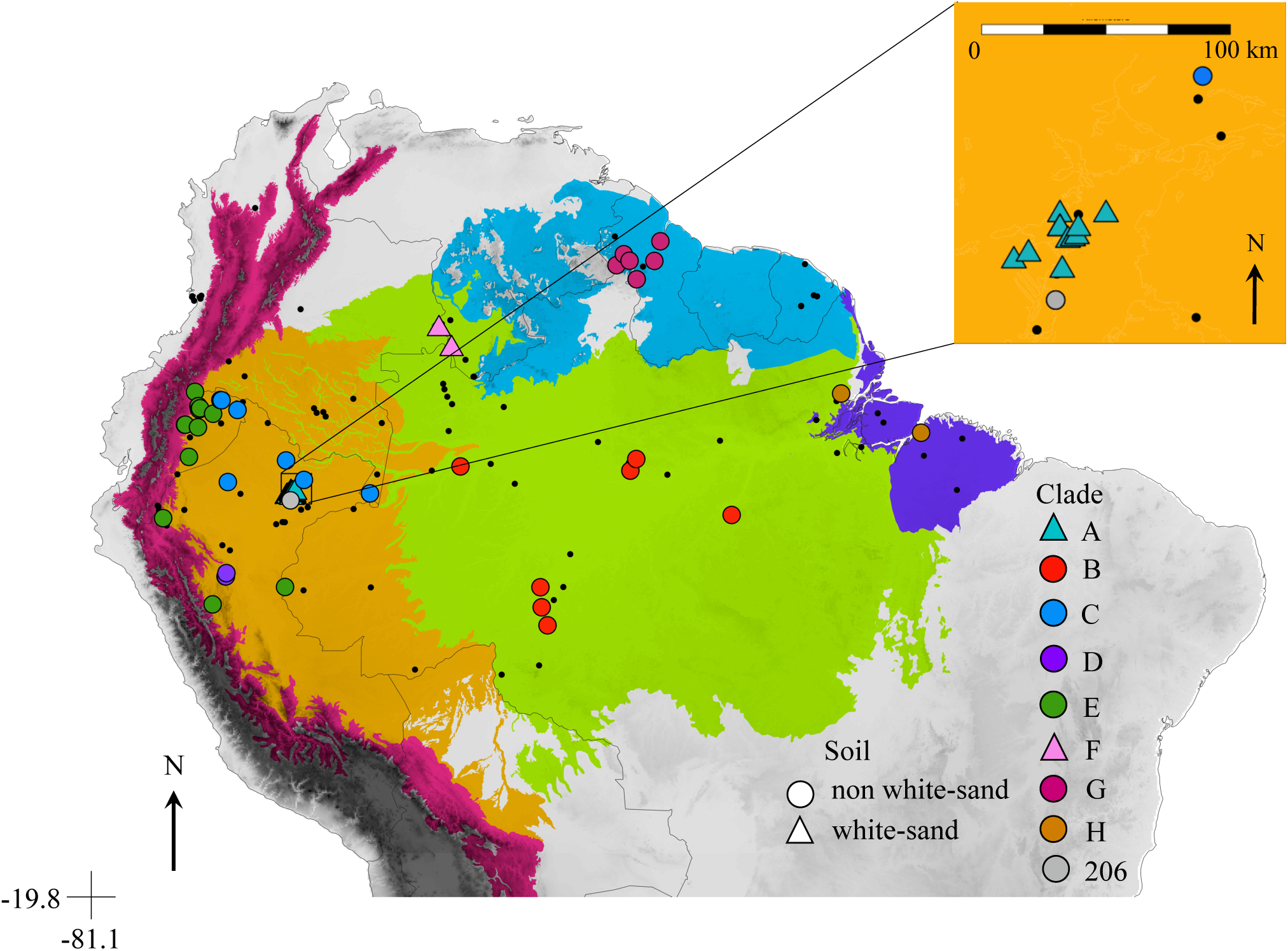
Range of *Protium subserratum*. Black points correspond to all herbarium accessions attempted for inclusion in this study. Colored points correspond to herbarium accessions with molecular data in this study colored by clade in Figure 2. Colored regions correspond with broad ecoregions. Blue = Guyana Shield Forests, Green = Amazon Forests, Orange = Western Amazon, Magenta = Andean Forests, Purple = Amazon River Estuary Forests.

### Bait Design

Custom Baits were designed utilizing three assembled transcriptomes: *Protium copal* (Schltdl. & Cham.) Engl. (Damasco et al., 2019), *Bursera simaruba* (L.) Sarg. and *Boswelia sacra* Flueck. (both downloaded through the 1KP project; onekp.com; [accessed 15 March 2019]; Carpenter et al., 2019) as well as the *Citrus sinensis* (L.) Osbeck draft genome, which was accessed from the Citrus Genome Database (citrusgenomedb.org; [accessed 15 March 2019]; Wu et al., 2014). Detailed methods are provided in the Supplementary Materials.

We utilized 1,841low or single copy loci (LSCL) gene sequences derived from the *P. copal* transcriptome. Target lengths ranged from 101 bp to 5,676 bp with an average length of 965 bp and were synthesized as part of a Custom MYbaits kit by Daicel Arbor Biosciences (Ann Arbor, MI, USA) in which baits were 80 bp in length with 3× tiling and 27 bp spacing.

### DNA extraction and library preparation

DNA was extracted from leaf material using an SDS and salt extraction protocol (Edwards et al. 1991). Extracted gDNA was eluted in 1x TE buffer and cleaned and concentrated with SPRI beads (AMPure XP; Beckman Coulter Life Sciences, Indianapolis, IN, USA). Sequencing libraries were prepared for the 183 samples with the highest quantities of DNA. Libraries were prepared using NEBNext FFPE DNA Repair Mix in combination with the NEBNext Ultra II DNA Library Prep kits for Illumina (New England Biolabs, Ipswich, MA, USA) per manufacturer recommendations with custom inline barcodes as described in Moore et al. (2018). All of our DNA template was highly degraded and thus fragmentation was not required.

### Targeted enrichment and sequencing

Libraries were multiplexed (4 - 24 samples per pool) for hybridization with each equimolar pool containing a total of 87 to 1000ng DNA. Outgroup taxa were pooled separately when possible. Each pool was divided into two or three separate reactions for enrichment using a Custom MYBaits target capture kit following the manufacturer’s protocol with slight modifications. Briefly, each reaction was incubated at 60°C for 40 hours, after which enriched products were combined. Block A blocking oligos provided in the MYBaits target capture kit were replaced with 1μl xGen Universal Blocking Oligo - TS-p7(6nt) and 1μl xGen Universal Blocking Oligo - TS-p5 (Integrated DNA Technologies, Coralville, IA, USA). The volume of water added to the reaction was reduced by 1.5μl to maintain a 20μl reaction volume. The Block C and Block O blocking oligos were replaced with an equivalent volume of SeqCap EZ Developer Reagent (Roche Diagnostics, Mannheim, Germany). Each reaction was amplified for 16 cycles. Final libraries were pooled in equimolar quantities and sequenced on two lanes of Illumina Nextseq500 75bp SR at the Oklahoma Medical Research Foundation Clinical Genomics Center (Oklahoma City, OK, USA).

### Data processing for phylogenomics

Demultiplexed sequences were trimmed to remove adapter sequences and quality filtered using TRIMMOMATIC (Bolger et al., 2014) with the settings (HEADCROP: 4, 5 or 6 depending on the length of the adapter sequences; SLIDINGWINDOW:5:20 and MINLEN:30). Sequence quality was examined with FastQC v0.11.7 (Andrews, 2010) before and after trimming to ensure the complete removal of adapters and low-quality sequences. Any individual with fewer than 3×10^5^ sequences were removed from the dataset.

We used the pipeline HybPiper v1.2 (Johnson et al., 2016) to process our cleaned data (Misiewicz, 2021). Briefly, reads were mapped to targets using BWA v0.7.12 (Li and Durbin, 2009) and then assembled into contigs using SPAdes v3.11.1 (Bankevich et al., 2012; Slater & Birney, 2005) with a coverage cutoff of four. Assembled contigs were then aligned with their associated target sequence using Exonerate (Slater and Birney, 2005). We used the “intronerate” function to retrieve off-target sequences as well as exons (jointly referred to as “supercontigs”). We identified potential paralogs for removal in our dataset using paralog_investigator.py. We further filtered our dataset to remove any locus with less than 25% of the average coverage length for the exon, loci present in less than 75% of the samples and individuals missing more than 70% of the target sequences. We initially inferred trees using both exons and supercontigs. The backbone topology was similar for both datasets, however internal nodes near the tips lacked support with only the exon data (data not shown). Conversely, while the inclusion of off-target sequences improved node support near the tips, off-target sequences were not recovered for all individuals. Thus, our supercontig alignments were often missing significantly more taxa as compared to the exon only dataset. Therefore, in order to improve node support while maximizing the inclusion of taxa we constructed a final dataset that combined supercontigs, when available, with exons for individuals from which off-target sequences were not recovered. This dataset is hereafter referred to as superexons. To account for mis-assembled sequences, we further filtered our dataset to remove nucleotide positions with more than 50% missing data using TrimAl (Capella-Gutierrez et al., 2009) and then implemented Spruceup (Borowiec, 2019) to identify and remove outlier sequences using default parameters (window size = 20bp, overlap = 15bp) and a cutoff of 0.95. AMAS v1.0 (M. L. Borowiec, 2016) was used to split the final processed/trimmed alignment back into single-locus alignments and to generate summary statistics for the final concatenated alignment.

### Phylogenetic Inference

Maximum likelihood trees were constructed for each of the 1191 loci and the concatenated dataset. Individual gene trees were first reconstructed using RAxML v8.2.9 (Stamatakis, 2014) with the “-f a” option, GTRCAT model and 100 bootstrap replicates. Because sequence errors that may have been missed in earlier filtering steps often appear as long branches during phylogenetic reconstruction, we implemented TreeShrink v1.3.1 (Mai & Mirarab, 2018) using a false-positive tolerance rate of 0.05 to detect samples with unexpectedly long branches in each of our gene trees. Flagged samples were removed from their respective gene alignments. Finally, one individual, 206, was consistently identified with strong support as a hybrid in our hybrid detection analyses and was removed from the dataset. Individual gene alignments were concatenated using phyx (Brown et al., 2017) and RAxML was then used as described above to re-estimate individual gene trees as well as to infer the phylogeny using the full concatenated dataset.

The species phylogeny was estimated using ASTRAL-II v4.10.12 (Mirarab & Warnow, 2015) which uses relationships between quartets of taxa to estimate the overall species tree. The *ape* package in R was used to collapse nodes with <33% bootstrap support in our maximum likelihood gene trees in order to minimize the impact of gene-tree estimation error (Mirarab & Warnow, 2015; Sayyari & Mirarab, 2016). ASTRAL-II identified the species tree that shares the maximum number of quartet trees with the 1191 gene trees estimated with RAxML and local posterior probability support values were calculated. To assess conflict among our gene trees and the species tree, discordance between gene trees was calculated using PhyParts v0.0.1 (Smith et al., 2015) (https://bitbucket.org/blackrim/phyparts) and incongruence between gene trees for each node was visualized by mapping results onto the ASTRAL species tree using PhyPartsPieCharts v1.0 (https://github.com/mossmatters/MJPythonNotebooks).

### SNP Datasets

We used SeCaPr v1.1.4 (Andermann et al., 2018) to generate a psuedoreference with the consensus sequences for each target locus to call SNPs. We then mapped our cleaned, paired reads to this pseudoreference using BWA v0.7.12 (Li & Durbin, 2009). Duplicates were removed, and SNPs were called using HaplotypeCaller, part of the Genome Analysis Toolkit (GATK) v4.0.1.2(McKenna et al., 2010). We created eight SNP datasets in total (Table 1). The first dataset included all *P. subserratum* individuals from clade A-H and one *P. ferrugineum* outgroup individual (SNP1). In order to assess population structure, we created an additional seven datasets to account for different groupings based on hierarchical levels observed in our phylogeny as well as to investigate potential gene flow among lineages in close geographic proximity to one another. Only one individual from clade F passed filtering and was thus omitted from these seven SNP datasets. SNP datasets for structure analysis included all individuals from clades A-H (but excluding clade F) (SNP2), all individuals from clades A-E (SNP3), all individuals from clades D and E (SNP4), all individuals from clades A and B (SNP5), all individuals from clades G and H (SNP6), all individuals from clades A and C (SNP7) and all individuals from clade B and C (SNP8). The two individuals with ambiguous placement in the phylogeny (114 and 052) were grouped with individuals from clade D and the individual consistently identified as a hybrid (206) was grouped with individuals from clade C. SNPs were combined for all individuals in a dataset using GATK CombineGVCFs and genotypes were jointly called across each dataset using GATK GenotypeGVCFs. For each dataset we used VCFtools (Danecek et al., 2011) to remove indels as well as SNPs within 5 bases of an indel. BCFtools v1.8 (Li, 2011) was used to apply the following SNP filtering thresholds: mapping quality (>40%), depth (>25), quality by depth (>2), minimum depth across individuals (>10). We also removed SNPs with a minor allele frequency <0.01, and variants with >20% missing genotypes. Only biallelic SNPs were retained and monomorphic sites were excluded. Finally, linked SNPS were removed with Plink (Purcell et al., 2007) by filtering the dataset to remove SNPs exceeding a variance inflation factor (VIF) using a window size of 50kb, a variant count of 5, and linkage threshold of 2 (--indep 50 5 2).

**Table 1.** SNP datasets

| SNP Dataset | Clades | #SNPS | # Genes | Analysis |
| --- | --- | --- | --- | --- |
| SNP1 | A, B, C, D, E, F, G, H, outgroup | 4,829 | 963 | Dsuite, PCA |
| SNP2 | A, B, C, D, E, G, H, 206 | 4199 | 918 | Structure |
| SNP3 | A, B, C, D, E, 206 | 3719 | 885 | Structure |
| SNP4 | C, D, E, 206 | 2229 | 783 | Structure |
| SNP5 | A, B, 206 | 2195 | 776 | Structure |
| SNP6 | G, H | 1364 | 712 | Structure |
| SNP7 | A, C, 206 | 1699 | 700 | Structure |
| SNP8 | B, C, 206 |  |  | Structure |

### Hybrid Detection

Under a neutral model of evolution, incomplete lineage sorting should lead to similar levels of shared ancestral polymorphisms among closely related lineages. However, in a scenario where hybridization has occurred, lineages experiencing gene flow will share a greater number of alleles relative to the background signal attributed to incomplete lineage sorting. Thus, by taking into account the allele distribution in three taxa relative to an outgroup (((P1,P2),P3,)Outgroup), site-pattern based frequency methods (i.e., ABBA-BABA tests) are able to distinguish hybridization from incomplete lineage sorting (Durand et al., 2011; Green et al., 2010). We used multiple site-pattern based frequency methods to infer hybridization across our datasets. First, we used HyDe (Blischak et al., 2018) to detect phylogenetic invariants for every combination of trios, for all individuals, using our concatenated superexon alignment with the run_hyde.py script. Next, the Dtrios function in Dsuite (Malinsky et al., 2021) was used to calculate Patterson’s *D* statistics and f4-ratio statistics using the full SNP dataset (SNP1) (Table 1). The significance of each test was computed using 100 jackknife resampling runs and Benjamini-Hochberg (BH) corrections were applied to account for false discovery.

### Substructure/Lineage Differentiation

Complex demographic, environmental and historical processes can influence the contemporary genetic structure of species resulting in different levels of organization (Meirmans, 2015). Furthermore, wide-scale population genetic structure can mask genetic differentiation among locally divergent populations (Pritchard & Wen 2004, Evanno et al., 2005; Pritchard et al., 2010). To account for these factors, we utilized a hierarchical approach to explore population structure across our dataset using Bayesian Markov chain Monte Carlo clustering analysis implemented in ParallelStructure (Besnier & Glover, 2013), an R-based implementation of STRUCTURE 2.3.3 (Pritchard et al., 2000) via the CIPRES Science Gateway (Miller et al., 2010). Structure analyses were run individually for each SNP dataset using the admixture model and assuming correlated allele frequencies. In order to estimate K, a burn-in period of 5 x 10^5^ generations was followed by 10^5^ Markov chain Monte Carlo generations for each value of K = 1 through K= 5-10 depending on the number of groups included in the analysis (Fig. S1-S8). Simulations were repeated twenty times for each value of K. Structure Harvester (Earl & vonHoldt, 2012) was used to interpret the output as described by Evanno et al. (2005) and Pritchard et al. (Pritchard et al., 2000). CLUMPP (Jakobsson & Rosenberg, 2007) and DISTRUCT 1.1 (Rosenberg, 2003) were implemented in Clumpak (Kopelman et al., 2015) to average admixture proportions over all runs and visualize the final. Principal components analysis (PCA) was carried out to identify differences in allele frequencies among populations and confirm our STRUCTURE results twice. The PCA was run twice, first analysis was with all *P. subserratum* individuals from the SNP1 dataset (outgroup individual removed), and our second analysis also used the SNP1 dataset, but excluded the outgroup individual and individuals from clades G and H. Analyses were carried out using the R package adegenet v2.1.1 (Jombart, 2008) to identify differences in allele frequencies among populations and confirm our STRUCTURE results. Finally, genetic differentiation among all lineages was assessed on SNP dataset SNP2 (Table 1) using pairwise comparisons of F_ST_ calculated in GenoDive v3.03 (Meirmans, 2020) generating confidence intervals over 10^5^ permutations.

### Georeferencing and Environmental Associations

*Protium subserratum* specimens that lacked geographic coordinates were georeferenced using U.C., Berkeley’s BerkeleyMapper (http://berkeleymapper.berkeley.edu/).

White-sand soils can have patchy distributions at fine scales, particularly in the western Amazon. Thus, we aimed to identify which of our specimens were associated with white-sand soils while also avoiding mis-classification due to uncertainty associated with geo-referencing by using a two-step process. First, we mapped only specimens that included geographic coordinates in their label data onto the most comprehensive map of known Amazon white-sand forests to date (Adeney et al., 2016). Then, the specimens that we georeferenced were assessed and considered to be white-sand inhabitants if they met two criteria: 1) Phylogenetic analysis placed them in the same as individuals with coordinates that mapped to known white-sand forests and 2) their georeferenced location was within 5km of known white-sand forests.

Similarly, because elevation can change significantly over short distances, particularly in the Andes, we only calculated elevation for specimens with known coordinates, thus, reducing error due to uncertainty associated with georeferenced specimens. Elevation was determined using Google Earth Pro v7.3 (Google, Mountain View, CA, USA).

### Ancestral Soil Habitat Estimation

To investigate the extent to which specialization onto white-sand habitats has occurred within *P. subserratum*, we estimated ancestral habitat affiliations. Using the ASTRAL phylogeny pruned to one tip per clade, we utilized BioGeoBEARS (Matzke, 2013) as implemented in rasp 4.0 (Yu et al., 2020) to test three different biogeographical models (BayAreaLIKE, DEC, and DIVALIKE) with and without jump dispersal (+j) which allows for founder event speciation. Soil habitats were assigned as either white-sand or non-white-sand. While we were unable to obtain usable sequence data for the true sister species to *P. subserratum*, previous research suggests that they are restricted to white-sand habitats (Daly and Fine, 2011; Fine et al., 2014). Thus, we designated our outgroup as a white-sand specialist. Lineages were allowed to simultaneously occur in both white-sand and non-white-sand habitats. The best-fit biogeographical model for each approach was identified using the Akaike information criterion corrected for sample size (AICc).

### Characterization of phenotypic variation

In order to investigate the extent to which phenotype is conserved within lineages and soil habitats, we characterized variation in five leaf morphological traits measured from specimens included in our phylogenetic dataset. Percent pubescence on the abaxial side of the leaflet blade and leaflet midrib, leaflet length, leaflet thickness, and the number of margin serrations per leaflet were collected from three individual leaflets per specimen and used to calculate the mean. Serration density was calculated by dividing the mean number of margin serrations per leaflet by the length of the leaflet. Pubescence percentage coverage on the abaxial side of the leaflet blade and pubescence percentage coverage on the leaflet midrib were visually estimated using a dissecting microscope to the nearest 10% within a haphazardly placed three by three-millimeter square. Three measurements were taken per leaflet on three leaflets per specimen and used to calculate mean. Morphological differentiation was assessed among individuals grouped by clade and soil type (white-sand or non-white-sand). To determine whether results from previous work that identifies morphological differentiation among white-sand and non-white-sand individuals from Peru holds with sampling across the range, we assessed differentiation across the full dataset. We conducted principal components analysis (PCA) to determine whether or not morphology is partitioned by soil type or clade. Kruskall Wallace tests were used to quantify the extent to which individual traits varied by clade and soil type. Analyses were conducted in R version 3.6.3 (R Core Team 2020) in RStudio v1.4.1717 (R Studio Team 2020).

## RESULTS

### Target Enrichment

Our final dataset included 1,191 loci for 67 individuals. Forty-seven loci were removed from the dataset because they were flagged as potentially paralogous and another 603 loci were filtered due to low representation across individuals. Out of 183 sequenced individuals 117 were dropped due to low sequencing coverage. Off-target sequences were recovered from an average of 43 of the final 67 individuals per gene (range 26 - 67). The final dataset included 59 *P. subserratum* accessions and eight outgroup accessions (seven *P. ferrugineum* and one *P. amazonicum*). The total alignment for the 1,191-locus supermatrix was 1,122,006 bp in length and with a total of 75,174,402 base pairs with 63,028 variable sites, 21,875 parsimony-informative sites, and 24% missing data. After cleaning and filtering, our SNP calling approach for all datasets yielded between 4,829 - 1,364 SNPs from 712 - 963 genes per individual (Table 1).

### Phylogenetic Inference

A hybrid individual (see *Hybrid Detection*) was excluded from the alignment, leaving 65 individuals for inclusion in our final phylogenetic dataset. Topologies from the concatenated RAxML and coalescent-based ASTRAL analyses were highly congruent and produced well-resolved phylogenies with the majority of differences corresponding to intraclade relationships, which are unlikely to be well represented by bifurcating trees (Fig. 2). Most major clades received definitive bootstrap support (BS=100%) in the concatenated maximum likelihood phylogeny and strong local posterior probability support (>0.9) in the Astral species tree (Fig 2). Both phylogenies resolved 8 major clades, 7 of which had strong support in both trees (clades A, C, D, E, F, G, H) (Fig 2). Two individuals, 052 and 114 could not be placed with confidence in either phylogeny (Fig. 2). Gene tree concordance was highest at the deepest nodes, supporting the monophyly of *P. subserratum* as well as the split between *P. subserratum* clades in the Western and Central Amazon (clades A, B, C, D, E, F) and those in Guyana and Eastern Brazil (Clades G and H) (Fig. 2). Correspondingly, gene tree concordance was virtually non-existent for the most recently derived clades (A, B, C, D, E and F), even though they had high bootstrap and local posterior probability support, where increased sharing of ancestral alleles and ongoing gene flow are more likely occurring (Fig. 2).

**Figure 2:**
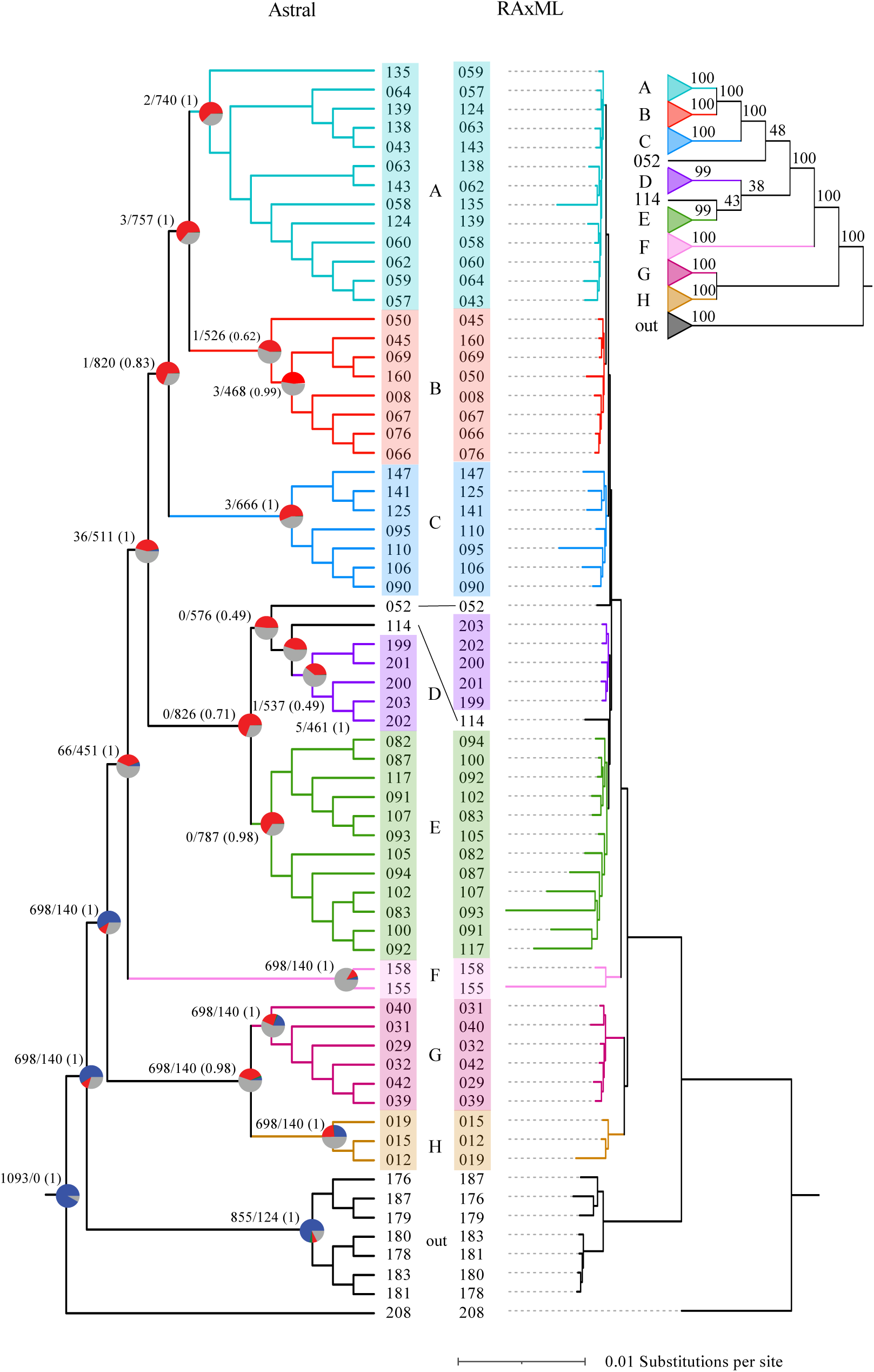
Astral species tree (left) and RAxML tree (right) with branch lengths and clade names in between trees. Geography and habitat represented by each clade is as follows: clade A: Peru, lowland, white-sand; clade C: Peru, lowland, non-white-sand; clade D: Peru, midelevation, non-white-sand; clade E: Peru, non-white-sand, montane; clade F: Venezuela, white-sand; clade G: Guyana, non-white sand; clade H: Eastern Brazil, non-white-sand. On the Astral tree, pie charts at nodes represent gene tree discordance at each node. Blue = proportion of trees concordant with the topology at that node, Red = proportion of trees with discordant topologies at that node, grey = proportion of trees with unsupported topologies at that node, and green = proportion of trees with a second alternative topology. Above each node, the number of concordant trees over the number of conflicting trees for that node. Astral support values are in parentheses. For the RAxML tree support values for each node are depicted in the cartoon inset. Lines between the two phylogenies indicate clade level disagreement between the Astral and RAxML topologies.

### Hybrid Detection

Using SNP data, a total of 16,215 tests were performed using Dsuite, of which 14 were significant (P-value < 0.05 after Bonferroni correction) (Table 2). D-statistics for significant combinations ranged from 0.38 to 0.22. Individual 206 was identified in 71% of statistically significant combinations, sharing alleles from individuals found in each of the five major clades from the Western and Central Amazon (clades A-E). Additional combinations with shared alleles included individual 090 (clade C) and individual 045 (clade B), individual 058 (clade A) and individual 203 (clade D), individual 90 (clade C) and individual 076 (clade B), and individual 057 (clade A) and individual 199 (clade D) (Table 2).

**Table 2:**
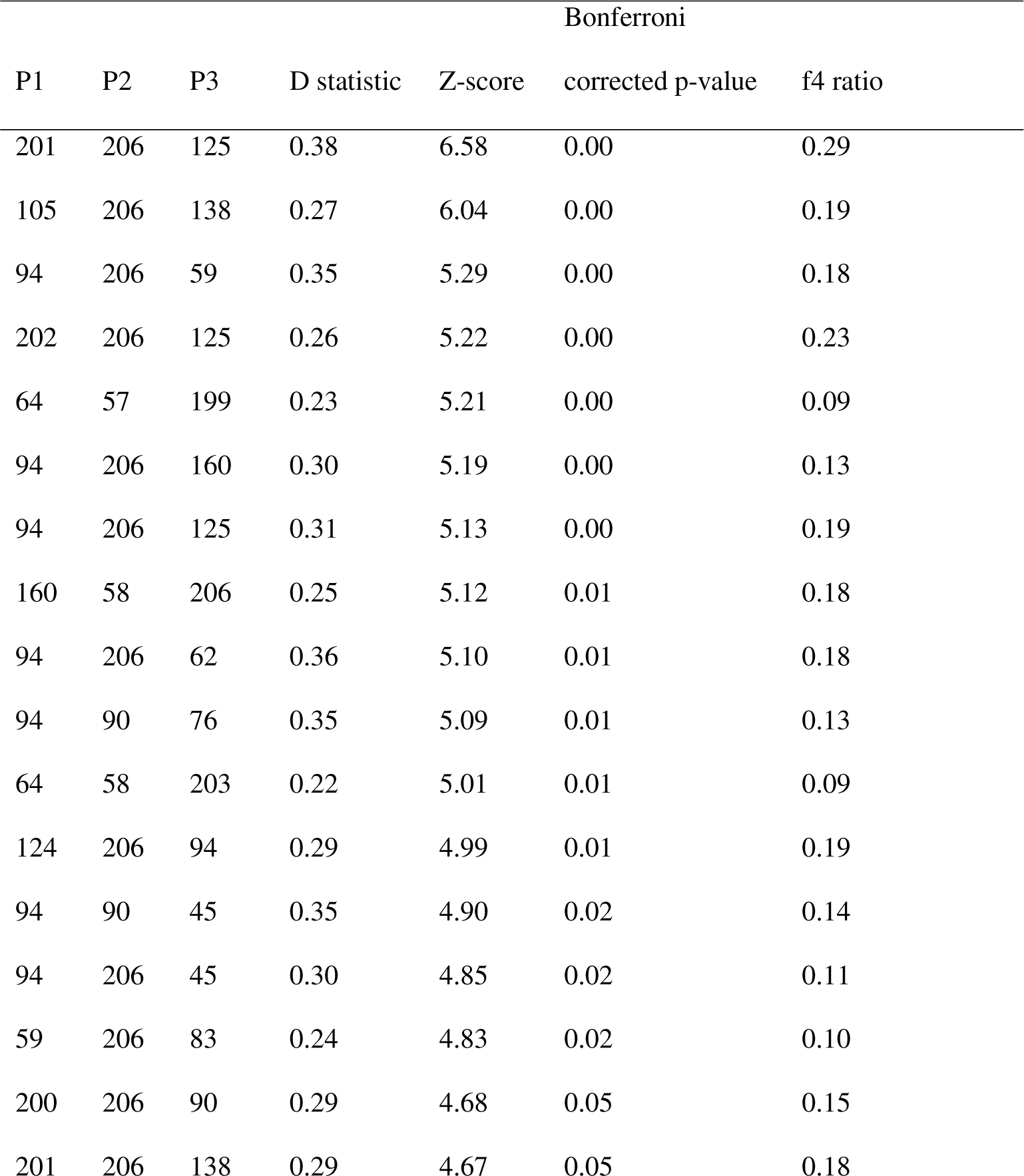
Dsuite results

| P1 | P2 | P3 | D statistic | Z-score | Bonferroni |  |
| --- | --- | --- | --- | --- | --- | --- |
|  |  |  |  |  | corrected p-value | f4 ratio |
| 201 | 206 | 125 | 0.38 | 6.58 | 0.00 | 0.29 |
| 105 | 206 | 138 | 0.27 | 6.04 | 0.00 | 0.19 |
| 94 | 206 | 59 | 0.35 | 5.29 | 0.00 | 0.18 |
| 202 | 206 | 125 | 0.26 | 5.22 | 0.00 | 0.23 |
| 64 | 57 | 199 | 0.23 | 5.21 | 0.00 | 0.09 |
| 94 | 206 | 160 | 0.30 | 5.19 | 0.00 | 0.13 |
| 94 | 206 | 125 | 0.31 | 5.13 | 0.00 | 0.19 |
| 160 | 58 | 206 | 0.25 | 5.12 | 0.01 | 0.18 |
| 94 | 206 | 62 | 0.36 | 5.10 | 0.01 | 0.18 |
| 94 | 90 | 76 | 0.35 | 5.09 | 0.01 | 0.13 |
| 64 | 58 | 203 | 0.22 | 5.01 | 0.01 | 0.09 |
| 124 | 206 | 94 | 0.29 | 4.99 | 0.01 | 0.19 |
| 94 | 90 | 45 | 0.35 | 4.90 | 0.02 | 0.14 |
| 94 | 206 | 45 | 0.30 | 4.85 | 0.02 | 0.11 |
| 59 | 206 | 83 | 0.24 | 4.83 | 0.02 | 0.10 |
| 200 | 206 | 90 | 0.29 | 4.68 | 0.05 | 0.15 |
| 201 | 206 | 138 | 0.29 | 4.67 | 0.05 | 0.18 |

We also tested for hybrids with our sequence data using HyDe. In total 97,528 tests were performed. In addition to the full results table, HyDe also reports the subset of tests deemed to be significant and producing reasonable gamma values. Seventy-five of our tests fell into this subset with 12 individuals identified as hybrids (Table 3). The mean Z-score across all significant tests was 5.15 (range: 4.88-6.01 and the mean gamma value was 0.49 (range 0.11 to 0.75). Similar to our Dsuite results, individual 206 was identified as a hybrid in 45% of the significant results with genetic contributions from individuals found across each of the five major clades from the Western and Central Amazon (clades A-E). Individual 052 was detected as a hybrid in 16% of tests and individual 090 (clade C) was detected in 13% of tests. The remaining nine individuals were identified as hybrids in 6% or fewer of the tests. Individual 206 and 090 were the only individuals identified as hybrids by both methods.

**Table 3:** HyDe results

| P1 | Hybrid | P2 | Z-score | p-value | Gamma |
| --- | --- | --- | --- | --- | --- |
| 114 | 52 | 58 | 4.90 | 0.00 | 0.43 |
| 114 | 52 | 62 | 5.17 | 0.00 | 0.43 |
| 138 | 52 | 203 | 4.90 | 0.00 | 0.45 |
| 114 | 52 | 66 | 5.08 | 0.00 | 0.45 |
| 114 | 52 | 60 | 5.19 | 0.00 | 0.46 |
| 139 | 52 | 203 | 5.01 | 0.00 | 0.48 |
| 141 | 52 | 199 | 4.91 | 0.00 | 0.50 |
| 43 | 52 | 201 | 4.94 | 0.00 | 0.52 |
| 203 | 52 | 143 | 4.99 | 0.00 | 0.54 |
| 203 | 52 | 43 | 4.99 | 0.00 | 0.55 |
| 203 | 52 | 62 | 4.93 | 0.00 | 0.57 |
| 203 | 52 | 57 | 4.96 | 0.00 | 0.58 |
| 206 | 57 | 66 | 4.89 | 0.00 | 0.46 |
| 206 | 57 | 67 | 5.05 | 0.00 | 0.46 |
| 76 | 57 | 206 | 5.24 | 0.00 | 0.56 |
| 160 | 57 | 105 | 5.08 | 0.00 | 0.71 |
| 160 | 58 | 105 | 4.94 | 0.00 | 0.69 |
| 206 | 62 | 8 | 5.14 | 0.00 | 0.50 |
| 160 | 62 | 105 | 4.89 | 0.00 | 0.73 |
| 32 | 69 | 95 | 4.94 | 0.00 | 0.12 |
| 76 | 90 | 83 | 5.73 | 0.00 | 0.49 |
| 83 | 90 | 8 | 5.08 | 0.00 | 0.49 |
| 160 | 90 | 105 | 5.66 | 0.00 | 0.49 |
| 8 | 90 | 94 | 5.42 | 0.00 | 0.51 |
| 76 | 90 | 94 | 5.27 | 0.00 | 0.51 |
| 160 | 90 | 94 | 5.86 | 0.00 | 0.51 |
| 160 | 90 | 107 | 5.20 | 0.00 | 0.52 |
| 76 | 90 | 107 | 5.01 | 0.00 | 0.53 |
| 76 | 90 | 93 | 5.05 | 0.00 | 0.55 |
| 160 | 90 | 93 | 5.28 | 0.00 | 0.57 |
| 58 | 102 | 91 | 4.96 | 0.00 | 0.43 |
| 76 | 106 | 83 | 5.33 | 0.00 | 0.43 |
| 160 | 110 | 102 | 4.93 | 0.00 | 0.50 |
| 160 | 110 | 94 | 5.07 | 0.00 | 0.52 |
| 160 | 110 | 105 | 5.09 | 0.00 | 0.53 |
| 60 | 110 | 102 | 4.99 | 0.00 | 0.58 |
| 76 | 110 | 83 | 4.94 | 0.00 | 0.59 |
| 200 | 143 | 66 | 4.92 | 0.00 | 0.29 |
| 202 | 143 | 66 | 5.10 | 0.00 | 0.30 |
| 160 | 147 | 105 | 4.97 | 0.00 | 0.57 |
| 76 | 206 | 201 | 4.99 | 0.00 | 0.38 |
| 76 | 206 | 202 | 5.23 | 0.00 | 0.38 |
| 160 | 206 | 105 | 6.01 | 0.00 | 0.38 |
| 94 | 206 | 57 | 5.02 | 0.00 | 0.39 |
| 76 | 206 | 83 | 5.38 | 0.00 | 0.39 |
| 83 | 206 | 57 | 5.34 | 0.00 | 0.40 |
| 76 | 206 | 199 | 5.06 | 0.00 | 0.40 |
| 201 | 206 | 124 | 5.37 | 0.00 | 0.41 |
| 76 | 206 | 200 | 5.04 | 0.00 | 0.41 |
| 83 | 206 | 124 | 4.99 | 0.00 | 0.41 |
| 200 | 206 | 59 | 5.35 | 0.00 | 0.41 |
| 160 | 206 | 201 | 5.00 | 0.00 | 0.41 |
| 202 | 206 | 59 | 4.99 | 0.00 | 0.43 |
| 100 | 206 | 57 | 4.98 | 0.00 | 0.44 |
| 160 | 206 | 94 | 5.46 | 0.00 | 0.44 |
| 45 | 206 | 105 | 5.57 | 0.00 | 0.45 |
| 100 | 206 | 124 | 5.64 | 0.00 | 0.45 |
| 45 | 206 | 100 | 4.92 | 0.00 | 0.45 |
| 83 | 206 | 59 | 5.23 | 0.00 | 0.45 |
| 45 | 206 | 201 | 5.34 | 0.00 | 0.47 |
| 45 | 206 | 199 | 5.42 | 0.00 | 0.48 |
| 45 | 206 | 94 | 5.13 | 0.00 | 0.49 |
| 200 | 206 | 45 | 5.75 | 0.00 | 0.49 |
| 105 | 206 | 59 | 5.04 | 0.00 | 0.51 |
| 83 | 206 | 45 | 5.86 | 0.00 | 0.51 |
| 60 | 206 | 201 | 4.91 | 0.00 | 0.52 |
| 60 | 206 | 94 | 5.05 | 0.00 | 0.52 |
| 202 | 206 | 45 | 5.20 | 0.00 | 0.53 |
| 59 | 206 | 94 | 5.12 | 0.00 | 0.57 |
| 200 | 206 | 160 | 5.19 | 0.00 | 0.58 |
| 64 | 206 | 203 | 4.91 | 0.00 | 0.59 |
| 58 | 206 | 200 | 5.12 | 0.00 | 0.60 |
| 124 | 206 | 199 | 5.00 | 0.00 | 0.60 |
| 57 | 206 | 199 | 5.01 | 0.00 | 0.61 |

In order to assess the impact of putative hybrid individuals on the integrity of our tree, we reconstructed the phylogeny while systematically removing hybrid individuals. Only the removal of individual 206 significantly improved backbone support values (data not shown). Thus, we removed only individual 206 from the dataset for final phylogenetic analysis.

### Environmental Associations

Of 42 *P. subserratum* specimens with geographic coordinates, two mapped to known white-sand habitat documented by Adeney et al. (2016): Individual 124 (clade A), collected from the region of Loreto, Peru and individual 158 (clade F), collected from the state of Amazonas, Venezuela. An additional 13 georeferenced individuals in clade A, and one individual in clade F were classified as white-sand because they belonged to a clade containing a non-georeferenced individual that mapped to known white-sand forests and their georeferenced location was within 5km of known white-sand forests. All other individuals were considered to be non-white-sand (Fig. 1).

Clades were variable in the elevational niche they occupied (Fig. 3). Individuals from clades A, B, F, and H all occupied lowland areas in the Amazon. Clade G, which, in this study, is composed of individuals from Guyana, occupied higher elevations ranging from approximately 600 – 900 meters. We also observed that clades C, D, and E turn over across an elevation gradient extending from the lowland Amazon to tropical montane habitat, which is occupied by clade E.

**Figure 3.**
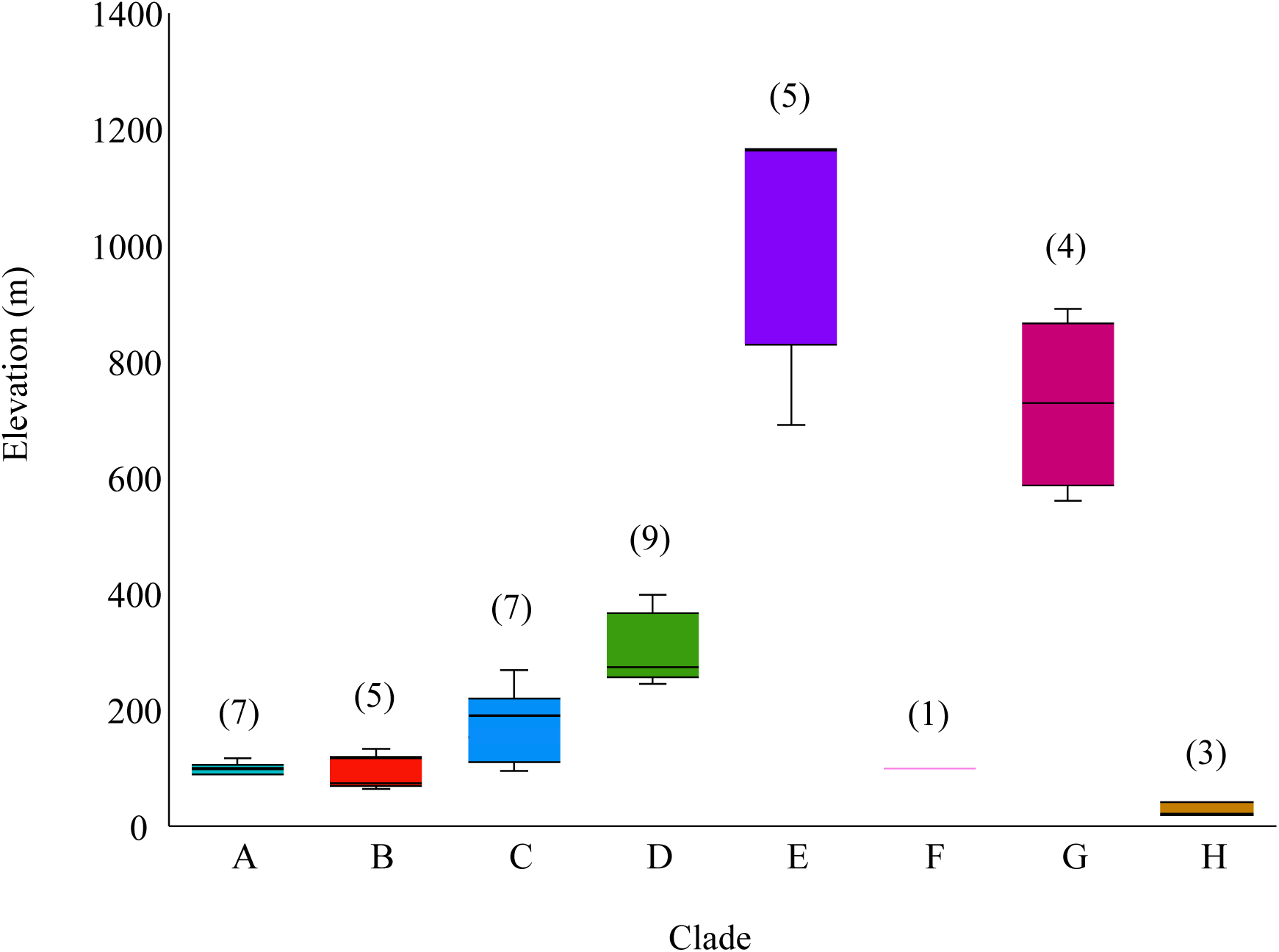
Box-whisker plots of elevation for individuals with coordinate data for each clade. Number of individuals per clade with coordinate data are listed above the box and whisker plot.

### Population Structure and Differentiation

Analysis of population structure revealed hierarchical structure. Using the full SNP dataset excluding outgroups and clade F (SNP2, Table 2), we found K=2 (Fig. 4, Fig. S1) to be the best supported model using the Evanno et al. (2005) method with lesser support for a secondary model observed at K=4 (Fig. 4, Fig. S1). Using the method of Pritchard et al. (2009), where the best model is inferred using the value of K with the greatest likelihood of the data (lnP(D)) or, if there is no clear maximum, the value of K at which lnP(D) begins to plateau, we found K=4 to be the best-supported model (Fig. S1). For K=2, individuals from clade G were assigned to one cluster, individuals from clades A-E were assigned to the second cluster, and individuals from clade H had significant contributions from both clusters (Fig. 4). For K=4, one genetic cluster corresponded with individuals from clade G, another was composed of individuals from clade H, a third cluster corresponded to individuals from both clade D and E and the final cluster corresponded with individuals from clades A and B (Fig 4). Individuals from clade C were represented as admixed between the clusters that corresponded to clades A and B and clades C and D (Fig. 4).

**Figure 4.**
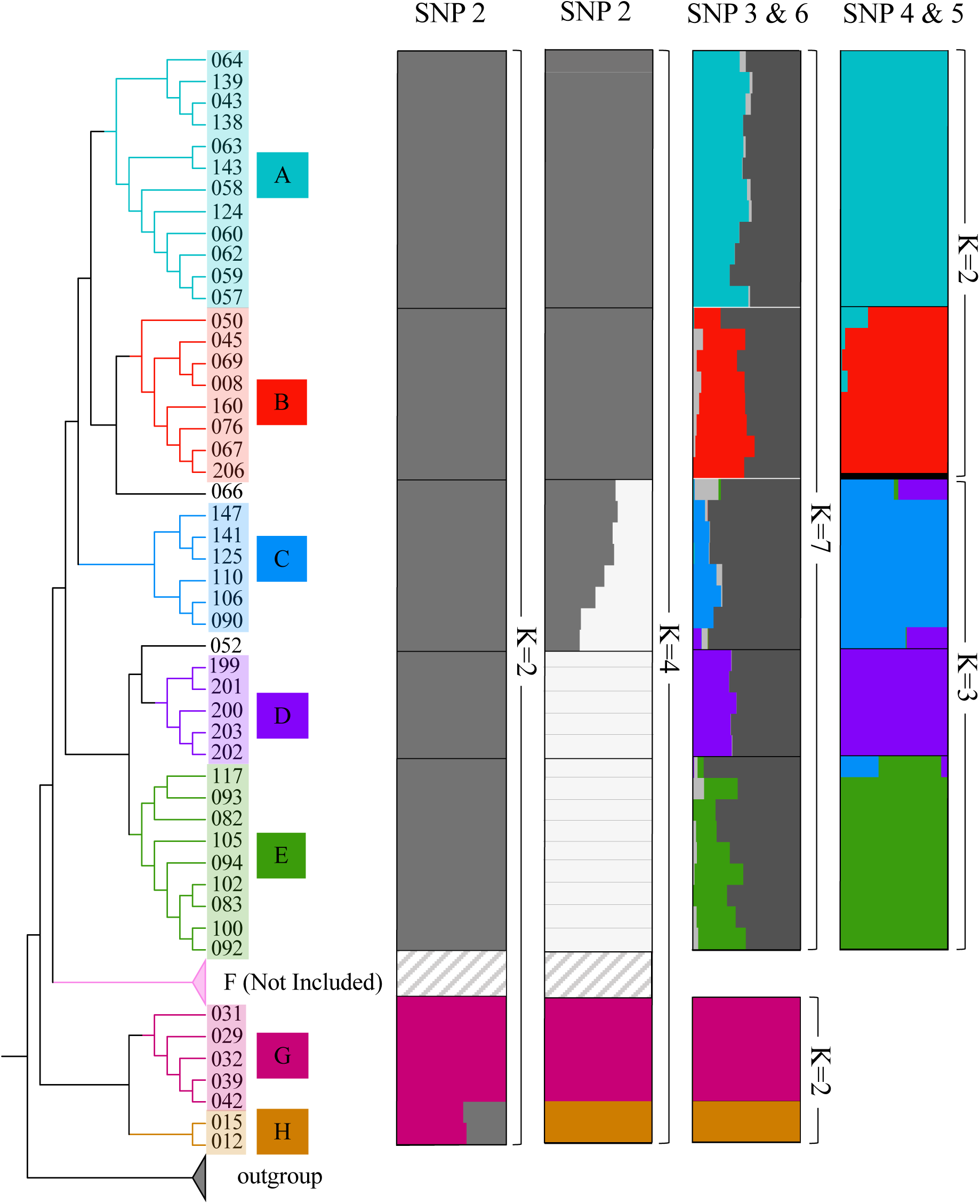
Best supported models from five Structure analyses using datasets SNP2, SNP3, SNP4, SNP5, and SNP6. With the exception of the hybrid individual, 206, which could not be placed with support, all individuals included in each analysis correspond to the Astral species tree.

Next, we analyzed various subsets of the data (Fig 4). For clades G-H (SNP6), K=2 was the best supported model for both methods, with genetic clusters corresponding to the two clades (Fig. S2). For clades A-E (SNP3), K=7 was favored with the Evanno method, although with low support, and there was no clear best K for the Pritchard method (Fig. S3). For K=7, five distinct groupings were observed, corresponding to the five clades, with the final two genetic groups appearing to represent genetic components shared across all clades. For clades A-B (SNP5), K=2 was the best supported model using both the Prichard and Evanno methods with each genetic cluster corresponding to the clade from which they were sampled (Fig S4). For clades C-E, (SNP4) K=3 was the best supported model using both methods and genetic groupings corresponded with each of the three clades (Fig. S5). Finally, while they do not represent monophyletic groups, we investigated the potential for admixture between clades A and C and clades B and C due to their close geographic proximity to one another (Fig. 1). For clades A and C (SNP7), we found K=4 to be the best supported model using the Evanno method and K=2 to be the best supported model using the Pritchard method (Fig. S6). However, both models depict two genetic populations corresponding to clades A and C (Fig. S8). In the K=4 model clade A and C each correspond to a genetic group with the remaining two genetic groups representing genetic components shared across individuals from both clades (Fig. S8). For clades B and C (SNP8), K=2 was the best supported model using both methods. Genetic clusters corresponded with each clade (Fig. S7, Fig S8).

Results from the PCA of SNP data were also consistent with groupings corresponding to the major clades identified within *P. subserratum* (Fig. 5). With all of the clades included, PC1 largely distinguished clades G and H from clades A-F (24.71% of the variation) (Fig 5A). When we excluded clades G and H, PC1 (15.49% of the variation) largely distinguished clades A-B and D-E from clades C and F (Fig. 5C). PC2 (9.09% of the variation) distinguished clades A and B (Fig. 5C), while PC3 (7.4% of the variation) differentiated clades D and E (Fig. 5D). Our pairwise F_ST_ values were consistently high between clade H and clades A, B, C, D, and E (Table 4). Within the western and central Amazon (clades A, B, C, D and E) F_ST_ values generally corresponded with geographic and phylogenetic distance (Table 4).

**Figure 5.**
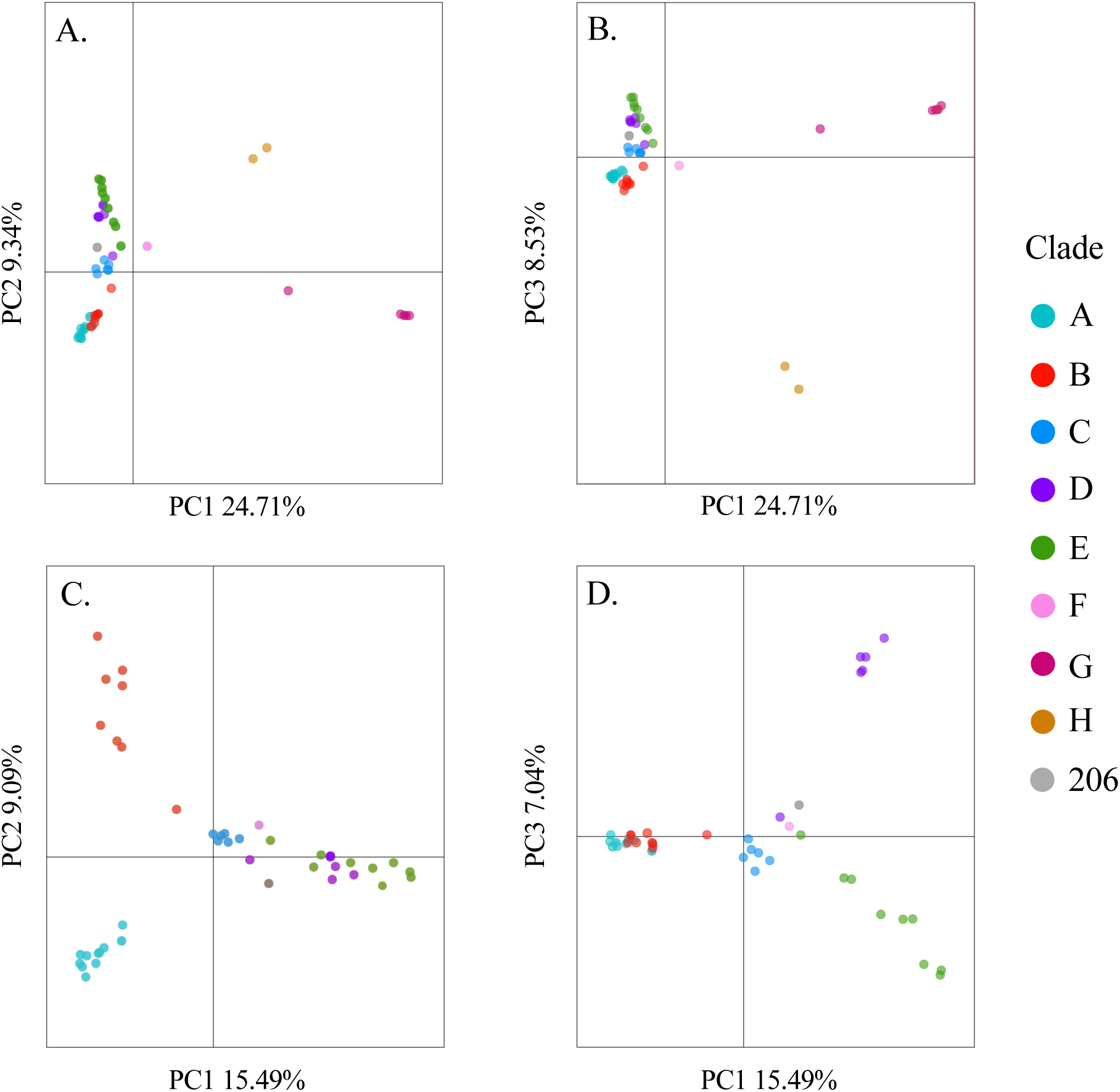
Principal components analysis of SNP1 data with points colored according to clade. Outgroup individuals are excluded in 5A and 5B. Outgroup individuals and individuals from clade G and H are excluded in 5C and 5D.

**Table 4.** Pairwise Fst values

| Table 4. Pairwise Fst values |  |  |  |  |  |  |
| --- | --- | --- | --- | --- | --- | --- |
| Clade | A | B | C | D | E | H |
| A | 0 |  |  |  |  |  |
| B | .204 | 0 |  |  |  |  |
| C | .198 | .195 | 0 |  |  |  |
| D | .272 | .268 | .215 | 0 |  |  |
| E | .288 | .291 | .201 | .191 | 0 |  |
| H | .643 | .636 | .666 | .634 | .640 | 0 |

### Ancestral Soil Habitat Estimation

The DIVA+j model was supported as the best model with the highest AIC score inferring two independent transitions from non-white-sand soil habitat to white-sand soil habitat. Without the +j parameter the best supported model was the DEC model. In this case the result was similar to the DIVA+j result except that ancestors at all nodes preceding a transition to white-sand were predicted to be soil generalists, occurring on both white and non-white-sand (Fig. 6).

**Figure 6:**
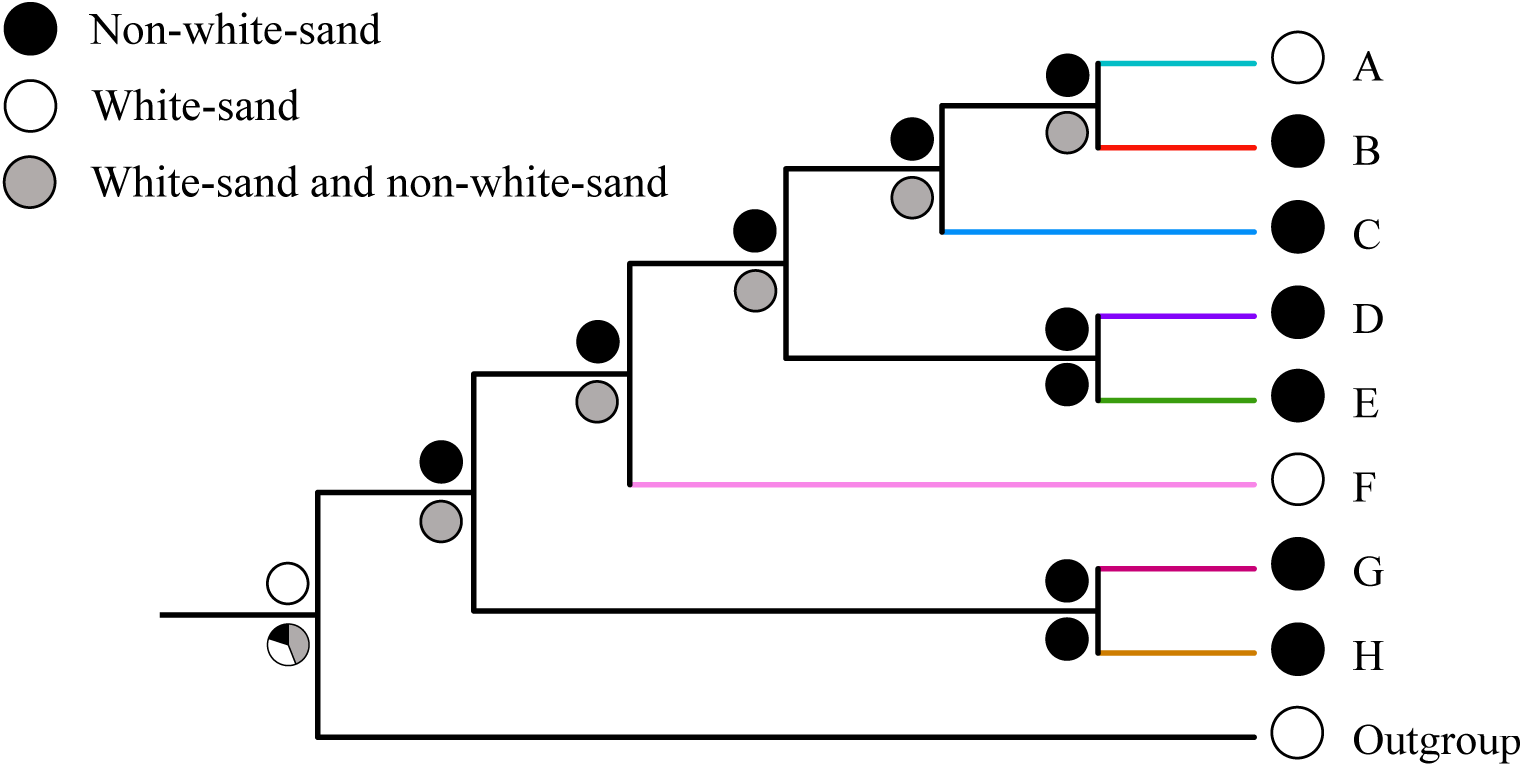
Results from the BioGeoBEARS analysis on the Astral species tree pruned to one individual per clade. Colored circles at the tips represent the soil that each clade inhabits. Results above the node correspond to the DIVA+j model and results below correspond to the DEC model. Results depicted are for soils predicted with >50% probability.

### Characterization of phenotypic variation

The first three principal components accounted for 81.8% of the variation in the data. However, leaf morphology was variable with no obvious partitioning by clade or soil (Fig. 7). When we examined how individual traits varied across groupings, we found significant differences by both clade and soil type (Table 5). When grouped, by clade we observed significant differences in serration density (Kruskal–Wallis χ ^2^ = 33.19, *P* < 0.001, Fig. S9C), pubescence on the abaxial side of the leaflet blade (%) (Kruskal–Wallis χ ^2^ = 14.44, *P* =0.04, Fig. S9E), average leaflet thickness (Kruskal Wallis χ ^2^ = 23.36, *P* = 0.001, Fig. S9G) and pubescence on the abaxial side of the leaflet midrib (%) (Kruskal–Wallis χ ^2^ = 17.71, *P* = 0.01, Fig. S9I). Significant differences in pubescence on the abaxial side of the leaflet blade (%) (Kruskal–Wallis χ ^2^ = 4.71, *P* = 0.03, Fig. S9F), average leaflet length (Kruskal–Wallis χ ^2^ = 4.09, *P* = 0.04, Fig. S9B), and the serration density (Kruskal–Wallis χ ^2^ = 20.42, *P* < 0.001, Fig. S9D) varied significantly by soil type. The remaining traits did not have significant variation among clades (average leaflet length, Fig. S9A) or soil types (average leaflet thickness, Fig. S9H; pubescence on the abaxial side of the leaflet midrib, Fig. S9J).

**Table 5.** Vegetative morphological traits by clade and soil (Kruskall-Wallis Test)

|  |  |  |  |  |  |  |
| --- | --- | --- | --- | --- | --- | --- |
| Morphological variable | Clade |  |  | Soil |  |  |
| | $\chi^2$ | p-value | df | $\chi^2$ | p-value | df |
| Average leaflet length | 9.90 | 0.19 | 7 | 4.10 | 0.04 | 1 |
| Serration density | 33.19 | 0 | 7 | 20.42 | 0 | 1 |
| Leaflet pubescence | 14.44 | 0.04 | 7 | 4.71 | 0.03 | 1 |
| Vein pubescence | 17.71 | 0.01 | 7 | 2.86 | 0.09 | 1 |
| Leaflet thickness | 23.36 | 0.001 | 7 | 0.06 | 0.81 | 1 |

## Discussion

*Protium subserratum* is a widespread and common tree found across the Amazon basin. In this study we utilized herbarium specimens to provide the most comprehensive range-wide sampling of the species to date. We found that genetic diversity within *P. subserratum* corresponded to at least eight independently evolving clades defined by geography and environment. Specialization onto white-sand soils evolved independently at least two times within the species complex. While previous studies have demonstrated strong correlations between phenotype and soil in Loreto, Peru, this association did not hold with sampling across the range owing to substantial variation in leaf morphology both within and among clades. Finally, cryptic diversity at the base of the Andes was found to be associated with elevational shifts.

### Evolutionary History of P. subserratum

Our results were largely in line with previous research but provided greater resolution and stronger support. Our study also included much broader sampling across the range which enabled us to identify six additional and previously undetected monophyletic clades. The two earliest diverging clades in our phylogeny (clade G and H) are sister to one another and correspond respectively to non-white-sand individuals from Guyana and non-white sand individuals from eastern Brazil. We also found that white-sand and non-white sand morphotypes formed paraphyletic groups. Specialization onto white-sand soils has occurred at least twice within the species (clades A and F) with white-sand clades found nested within non-white-sand clades. Non-white sand clades A and B are restricted to the western and central lowland Amazon while clades E and D are composed of non-white-sand individuals occupying different elevational ranges at the base of the Peruvian Andes. While we were able to identify a total of eight monophyletic clades defined by geography, soil and elevation, this may not represent the full extent of diversification within *P. subserratum.* Notable gaps in our sampling include the Colombian Caquetá (morphotype 4; Daly & Fine, 2011), and individuals growing on white-sand soils from Guyana.

Future work should aim to increase sampling near Iquitos, Peru, the only area where three soil specialist ecotypes are known exist in parapatry. Population genetic work by Misiewicz and Fine (2014) used microsatellite data to demonstrate that non-white-sand populations in Loreto, Peru, near Iquitos, were diverging across habitats with genetically and morphologically distinct populations found on brown-sand and clay soils. We did not detect distinct clades associated with brown-sand and clay soils in our dataset; however, this may be in part due to the fact we were unable to obtain quality genetic data for more than one non-white-sand specimen in the immediate region sampled by Misiewicz and Fine (2014). If the population divergence previously detected is restricted to the region sampled in Misiewicz and Fine (2014), then our sampling would have been insufficient to detect the same pattern. Alternatively, our exon capture data may not provide sufficient resolution to show the presumably very recent population level divergence observed in that study. Interestingly, the single clay soil individual in our study that was also sampled in Misiewicz and Fine (2014) (individual 206) was identified with confidence as a hybrid in our analyses with genetic contributions from white-sand clade A, and non-white-sand clades B and D. This individual was not identified as a recent hybrid in Misiewicz and Fine (2014), nor did it represent an intermediate phenotype. One possibility is that individuals representing the clay ecotype identified in Misiewicz and Fine (2014) are derived from a historic hybridization event. This could explain why we were able to detect a signal of hybridization using sequence data while microsatellite data, which is likely to reflect very recent population genetic dynamics, did not detect a signal of ongoing hybridization. Nonetheless, additional sampling is needed to clarify the origin and phylogenetic position of that group.

### Phenotypic Differentiation and Soil Type

Based on our examination of herbarium specimens, there is no single morphological character or combination of characters that are associated solely with white-sand specialists. We found that leaf morphology across white-sand individuals is more or less consistent with the description provided in Daly and Fine (2011) where white-sand individuals from both Peru and Venezuela (clade A and F) consistently exhibited entire leaflet margins (Fig. S9C). Additional characters such as the presence of leaflet pubescence and thickness that were identified by Misiewicz and Fine (2014) as being associated with Peruvian white-sand individuals were much more variable both within and across white-sand clades. For example, pubescence on the abaxial side of the leaflet blade and midrib was highly variable within the Peruvian white-sand clade (clade A) and largely absent from individuals in the Venezuelan white-sand clade (clade F) (Fig. S9E and S9H). Similarly, individuals from the Venezuelan white-sand clade (clade F) exhibited thicker leaflets than white-sand individuals from Peru (clade A) (Fig. S9G). However, as clade F only included two individuals, more sampling is needed to confirm differences among these clades.

While white-sand individuals consistently presented entire leaflet margins, identifying an individual’s edaphic affinity based on morphology alone was still challenging when considering specimens from across the range. This difficulty was due primarily to the high degree of morphological variation observed both within and across non-white-sand clades (clades B, C, D, E, G, H), which often overlapped with the white-sand phenotype. For instance, entire leaflet margins, a character consistently observed in individuals from white-sand (Fig. S9D) is also observed in some individuals from non-white-sand clades (Fig. S9C). Furthermore, with the exception of individuals from clade H (“Morphotype 1”), which are unique in that they consistently exhibit entire leaflets with impressed abaxial tertiary veins (Daly & Fine 2011) we were unable to identify characters useful in differentiating among non-white-sand clades.

The extent to which morphological traits observed in white-sand and non-white-sand individuals from across the range are due to plasticity is largely unknown. A two-year reciprocal transplant study demonstrated that morphological variation observed between seedlings associated with white-sand and non-white-sand habitats in Loreto, Peru was due to a combination of plasticity and genotype (Fine et al. 2013b). Leaf pubescence was determined to be genetically based, while leaflet thickness and toughness appeared to be plastic. No other morphological characters, including differences in leaflet serrations, were investigated.

### Lineage Diversification in Common and Widespread Amazonian Trees

Our finding that *P. subserratum* is made up of numerous independently evolving clades is in line with the small number of other recent studies investigating diversification in widespread Amazonian tree taxa (Damasco et al., 2021; Roncal et al., 2013, 2015; Scotti-Saintagne et al., 2013; Vicentini, 2016). While our study did not directly address the mechanisms that drove diversification within this group, the geographic distribution of our clades did not correspond with major geographic breaks or barriers and thus is inconsistent with a hypothesis of vicariant speciation. Instead, it is much more plausible that long distance dispersal into novel habitats and geographic areas was likely the catalyst for initial population isolation followed by lineage divergence. This hypothesis is in line with other studies of diversification of Amazonian taxa, including studies of within species diversification (Damasco et al., 2019; Dexter et al., 2017; Roncal et al., 2015; and discussed in Dick and Pennington, 2019). For instance, a phylogeographic study by Roncal et al. (2015) suggested that *Astrocaryum* sect. *Huicungo* (previously *Astrocaryum murumuru*) likely experienced repeated dispersals into a variety of distinct habitats in the western Amazon, which led to the establishment of fragmented populations that were sufficiently spatially isolated as to undergo diversification in spite of being separated by geographic barriers that were permeable to gene flow (Roncal et al. 2015). Damasco et al. (2021) investigated diversification within *Protium heptaphyllum* and found that recent long-distance dispersal into new geographic regions and habitats (including but not limited to white-sand habitats) has also likely played an important role in lineage diversification.

Parapatric ecological speciation across soil boundaries has been invoked as a potential mechanism driving diversification in *P. subserratum,* particularly in the Peruvian Amazon where genetically and phenotypically differentiated ecotypes are found adjacent to one another on white-sand and non-white-sand soils (Fine et al., 2013a; Misiewicz and Fine, 2014). Because this process is expected to yield the same phylogeographic signature as a scenario where lineage diverge occurs in isolation after a long-distance dispersal event into a novel habitat followed by secondary contact, our study and others (Damasco et al., 2021; Roncal et al., 2015) were unable to discern the role of natural selection in driving the initial divergence of ecologically contrasting lineages. Regardless of the initial mechanisms of reproductive isolation, specialization onto different habitats appears to have played an important role in lineage diversification within *P. subserratum* and other widespread taxa (Damasco et al., 2021; Roncal et al., 2015).

We detected little consistent signal of hybridization among clades. While our sampling did not allow us to explicitly test for introgression at contact zones between clades our results are bolstered by previous work in this system. Misiewicz et al. (2020) examined multiple barriers to reproduction between adjacent populations of soil specialist ecotypes found on white-sand and brown-sand in Loreto, Peru (clades A and B) and found that a combination of genetic and ecologically based barriers to reproduction acted to maintain near complete isolation. Similar to our study, the few recent studies examining lineage diversification within widespread Amazonian species did not specifically target contact zones. However, Damasco et al. (2021) detected some evidence of gene flow among their most recently diverged lineages.

## Conclusion

The results presented here suggest that *P. subserratum* represents a species complex that is in the processes of transitioning from a single widespread generalist species to multiple, range restricted, specialist species. Our work adds to the body of literature suggesting that widespread Amazonian species should not necessarily be assumed to represent genetically or ecologically uniform entities. Instead, Amazonian taxa that are considered to be atypical due to being common and widespread may actually include multiple lineages that are in the process of diversification.

## Supporting information

Supplemental Methods and Results

## Acknowledgements

We thank Charlotte Bony, John Mclaughlin, Charlotte Miranda, Larke Reeber, David Vu, and Tandena Wagner for their insightful comments on the manuscript. We also thank Lydia Smith and Chodon Sass for their invaluable advice on optimizing lab protocols for herbarium material, Austin Betancourt for assistance with data collection and Zachary Meyers for facilitating access to laboratory equipment. This work could not have been completed without access to specimens housed at the National Herbarium, the Missouri Botanic Garden and the New York Botanical Garden Herbarium. The computing for this project was performed at the OU Supercomputing Center for Education & Research (OSCER) at the University of Oklahoma (OU). In particular we would like to thank OSCER staff Patrick Calhoun, David Akin, and Horst Severini for their technical assistance throughout the project. This project was funded by a National Science Foundation Postdoctoral Research Fellowship in Biology to TMM, Award #1810989.

## Data Accessibility

Raw sequence data were deposited at the National Center for Biotechnology Information Short Read Archive (http://www.ncbi.nlm.nih.gov/sra/) database under the accession number BioProject ID PRJNA869083. Target sequences used in this study as well as the processed SNP data (STRUCTURE format), concatenated and aligned superexon data (fasta format), partition file for the concatenated data set, and Astral II and RAxML tree files are available from DataDryad (https://doi.org/10.6078/D1V41S).

## Author’s Contributions

TMM designed the study. TMM, TS and BC gathered and analyzed the data. TMM wrote the manuscript. AM and PF provided guidance on the study and commented on the manuscript.

