## Supplemental Methods and Results for "Parallel evolution and cryptic diversification in the common and widespread Amazonian tree, *Protium subserratum*"

### Supplementary Methods and Results

#### Bait design

Our bait set was designed to target single to low orthologous loci for phylogenetic reconstruction as well as genes in the terpene synthase and flavonoid gene families.

Secondary Defense Genes Involved in Herbivore Defense

Baits to target putative genes underlying secondary herbivore defense chemistry were designed from 287 transcripts from the *P. copal* transcriptome (Damasco et al., 2019) that were identified by Trinotate (Bryant et al., 2017) to be terpene synthase genes or genes involved in the flavonoid biosynthesis pathway.

Low to Single Orthologous Loci

Different bait design pipelines often identify non-overlapping loci (Vatanparast et al., 2018). Thus, to maximize the number of informative low to single orthologous loci (LSCN) we employed two methods. First, we used the *Marker Miner v1.0* (Chamala et al., 2015) pipeline, modified to include the reference genome of *Citrus sinensis.* LSCL loci identified by MarkerMiner were then further sub-selected using the *Goldfinder* python script (Vargas et al., 2019). Second, we identified LSCL loci using the AllMarker and BestMarker scripts described by (Kadlec et al., 2017).

*MarkerMiner*

Adding a new reference genome to the *Marker Miner* resources library requires protein-coding reference sequences containing masked introns, list of single copy genes, and a proteome blast database for the new reference taxa.

To create the protein-coding reference sequences containing masked introns we utilized the *Citrus sinensis* v1.1 genome and associated resources (Citrus Genome Database; citrusgenomedb.org) and manipulated them using BEDTools (Quinlan and Hall, 2010). Exons and genes were identified and extracted from the *C. sinensis* genome. Introns were isolated using the BEDTools subtract function to remove the exons from the full gene sequences. Identified introns were then used to create an intron masked genome using the BEDTools maskfasta function. To create the final database of intron masked reference sequences, gene sequences were extracted from the intron masked genome using the bedtools getfasta function.

To create a list of LSCL genes we utilized three assembled transcriptomes, *Protium copal* (Damasco et al. 2019), *Bursera simaruba* and *Boswelia sacra* (available through the 1KP project; onekp.com) and *C. sinensis* protein sequences (Citrus Genome Database; citrusgenomedb.org). Trinity ((Grabherr et al., 2011) was used to choose the longest isoform per gene from the *P. copal* transcriptome. The longest open reading frames from all three transcriptomes (1KP transcriptome downloads for *B. simaruba* and *B. sacra* and the longest isoform file for *P. copal*) were then identified using TransDecoder (Haas et al., 2013); https://github.com/TransDecoder). Orthologous protein sequences from all three transcriptomes and the *C. sinsensis* protein sequences were then identified using Orthofinder (Emms and Kelly, 2019, 2015). Finally, single gene orthogroups for *C. sinensis* were retained and compared against the intron masked reference to ensure single copy genes were present in both.

The *Citrus sinensis* proteome database was created using the BLAST makeblastdb function for *C. sinensis* protein sequences.

Once the new reference files were added to the MarkerMiner resource library transcripts of less than 700bp were filtered and queried to the *C. sinensis* proteome using reciprocal BLAST (Altschul, 1997; Altschul et al., 1990). BLAST results were retained if 70% or more of the transcript length aligned to the reference protein with at least 50% sequence similarity and if a minimum of 80% of the protein length was aligned to a transcript with at least 50% sequence similarity. Transcripts with top reciprocal BLAST hits against single copy genes were retained and classified as putative LCSL orthologs. LCSL orthologs were further sub-selected using the *Goldfinder* python script (Vargas et al., 2019) using the *C. sinensis* genome as a reference. Default arguments were used with the exception of bait length, which was set to 100 and bait coverage, which was set to 3. *Protium copal* sequences identified from this Goldfinder run were retained for the final bait-set. The analysis was then repeated using a more stringent similarity (75%) and the best *B. simaruba* sequences identified from this second run were chosen.

**AllMarkers.py and BestMarkers.py**

We used the AllMarker and BestMarker pipelines described by Kadlec et al. (2017) to identify additional putative orthologous loci using the *P. copal* and *B. simaruba* transcriptomes, and the *C. sinensis* genome. Markers were first selected using AllMarkers.py where transcriptomes are compared to identify homologues. Next, genes with multiple copies were identified for exclusion by using BLASTn to compare the transcriptomes against the reference genome. Remaining sequences were then filtered for at least 65% similarity. Optimal target sequences were selected using *BestMarkers.py.* using the following arguments; intron = yes, number of baits = 20000, intron length = 200, prioritize = length, bait length = 100, bait coverage = 300. *P. copal* sequences were chosen from this run. The analysis was repeated using higher similarity parameters (75%) and from this output *B. simaruba* sequences were chosen.

**Final Bait Set**

Loci identified across all runs for each pipeline were compared and duplicates were removed. The final bait set consists of 38,138 baits targeting 2,716 genes of which 2,430 are putative LSCL genes and 286 are putative chemistry genes. Across the total target set gene lengths vary between 13 and 5,676 bp with an average length of 973 bp. From the total set of targets 1,841 LSCL gene sequences were from the *P. copal* transcriptome and range from 101 bp to 5,676 bp with an average of 965 bp. 589 LSCL gene sequences derived from the *B. simaruba* transcriptome and ranged from 100 bp to 4,887 bp with an average length of 1,288 bp. Finally, 286 genes putatively from the terpene synthase and flavonoid biosynthesis pathway were derived from the *P. copal* transcriptome and ranged from 13 to 2,166bp with an average length of 380 Bp. The final bait set was synthesized at Arbor Biosciences (MYbaits http://www.arborbiosci.com/). Baits designed from the *P. copal* transcriptome were 80 bp long, with 3X tiling and 27 bp spacing. Baits from the *B. simaruba* transcriptome were 80 bp long, with 1X tiling and 80bp spacing.

Table S1: Table of all accessions sampled for this study

| Sample # | *Protium* Species | Georef-erenced? | Latitude | Longitude | Clade | Soil | Altitude (m) | Herbarium | Accession# | Collector | Collection # |
|---|---|---|---|---|---|---|---|---|---|---|---|
| 8 | *P. subserratum* | yes | -4.63641 | -55.95367 | B | non-ws | 85 | US | 3207949 | Prance | 25619 |
| 45 | *P. subserratum* | no | -9 | -63.25 | B | non-ws | 102 | MO | 5935108 | Thomas | 4991 |
| 50 | *P. subserratum* | no | -2.716667 | -66.7 | B | non-ws | 64 | MO | 6021774 | Daly | 4133 |
| 66 | *P. subserratum* | no | -2.883333 | -59.966667 | B | non-ws | 75 | NYBG | 1188635 | Lohmann | 69 |
| 67 | *P. subserratum* | no | -2.423889 | -59.733333 | B | non-ws | 134 | MO | 6016992 | Mori | 20565 |
| 69 | *P. subserratum* | yes | -8.2871 | -63.4802 | B | non-ws | 55 | NYBG | 2219383 | Melo | 620A |
| 76 | *P. subserratum* | no | -2.883333 | -59.966667 | B | non-ws | 75 | NYBG | 1188638 | Lohmann | 2888-08 |
| 160 | *P. subserratum* | yes | -7.49348 | -63.53034 | B | non-ws | 68 | US | 3101904 | Teixeira | 937 |
| 12 | *P. subserratum* | no | -1.383333 | -48.45 | H | non-ws | 21 | NYBG | 2219388 | Daly | 3759 |
| 15 | *P. subserratum* | no | 0.166667 | -51.616667 | H | non-ws | 42 | MO | 5998117 | Mori | 17489 |
| 19 | *P. subserratum* | no | -1.383333 | -48.45 | H | non-ws | 19 | NYBG | 1241780 | Daly | 3762 |
| 52 | *P. subserratum* | no | -7.483333 | -73.65 | E | non-ws | 249 | NYBG | 1241761 | Campbell | 8603 |
| 82 | *P. subserratum* | yes | -1.066933 | -77.618183 | E | non-ws | 434 | NYBG | 2408677 | Bensman | 99 |
| 83 | *P. subserratum* | no | 0.233333 | -77.216667 | E | non-ws | 350 | NYBG | 2408681 | Ceron | 9298 |
| 87 | *P. subserratum* | no | -1.166667 | -77.1 | E | non-ws | 384 | MO | 3800480 | Rubio | 218 |
| 91 | *P. subserratum* | yes | -0.46163 | -77.056008 | E | non-ws | 283 | NYBG | 2408684 | Jaramillo | 2173 |
| 92 | *P. subserratum* | no | -0.333333 | -77.083333 | E | non-ws | 274 | NYBG | 2408682 | Espinoza | 21 |
| 93 | *P. subserratum* | no | -1.066667 | -77.6 | E | non-ws | 398 | MO | 5296071 | Zuleta | 11 |
| 94 | *P. subserratum* | no | -0.416667 | -77.016667 | E | non-ws | 264 | NYBG | 2408694 | Palacios | 10766 |
| 100 | *P. subserratum* | no | -0.333333 | -77.083333 | E | non-ws | 274 | MO | 4227909 | Gudino | 78 |
| 102 | *P. subserratum* | no | -0.333333 | -77.083333 | E | non-ws | 274 | NYBG | 2408680 | Rubio | 288 |
| 105 | *P. subserratum* | no | -0.633333 | -76.5 | E | non-ws | 245 | US | 3466773 | Villa | 1582 |
| 107 | *P. subserratum* | yes | -2.33917 | -77.45793 | E | non-ws | 343 | US | 2406166 | Cazalet | 7672 |
| 114 | *P. subserratum* | yes | -8.171492 | -76.518763 | E | non-ws | 490 | NYBG | 2408695 | Vigo | 7264 |
| 117 | *P. subserratum* | yes | -4.76192 | -78.47969 | E | non-ws | 955 | MO | 5207501 | Diaz | 8386 |
| 29 | *P. subserratum* | no | 4.683333 | -59.716667 | G | non-ws | 891 | NYBG | 905015 | Clarke | 1161 |
| 31 | *P. subserratum* | yes | 6.18639 | -58.77734 | G | non-ws | 67 | MO | 5935429 | Mori | 8133 |
| 32 | *P. subserratum* | no | 5.416667 | -60 | G | non-ws | 791 | US | 3208491 | Pipoly | 11076 |
| 39 | *P. subserratum* | no | 5.240278 | -60.516111 | G | non-ws | 666 | NYBG | 2408699 | Clarke | 11414 |
| 40 | *P. subserratum* | yes | 5.4087 | -59.01656 | G | non-ws | 120 | US | 1743466 | Tutin | 231 |
| 42 | *P. subserratum* | no | 5.674722 | -60.226389 | G | non-ws | 561 | NYBG | 2449502 | Redden | 1587 |
| 206 | *P. subserratum* | no | -4.058778 | -73.432082 | C | non-ws | 120 | UC | | Misiewicz | TM28 |
| 43 | *P. subserratum* | yes | -3.84273 | -73.38752 | A | ws | 89 | MO | 4888592 | Rimachi | 4162 |
| 57 | *P. subserratum* | yes | -3.95052 | -73.40821 | A | ws | 143 | NYBG | 710229 | Fine | 288 |
| 58 | *P. subserratum* | no | -3.75 | -73.416667 | A | ws | 106 | NYBG | 2394340 | Vasquez | 3495 |

| | | | | | | | | | | | |
|---|---|---|---|---|---|---|---|---|---|---|---|
| 59 | *P. subserratum* | yes | -3.892134 | -73.530372 | A | ws | 101 | NYBG | 2394347 | Aronson | 1037 |
| 60 | *P. subserratum* | yes | -3.829083 | -73.355444 | A | ws | 92 | MO | 6205254 | Rimachi | 3088 |
| 62 | *P. subserratum* | no | -3.8 | -73.416667 | A | ws | 103 | NYBG | 2394334 | Vasquez | 13640 |
| 63 | *P. subserratum* | no | -3.75 | -73.25 | A | ws | 90 | MO | 3513641 | Vasquez | 2613 |
| 64 | *P. subserratum* | no | -3.8 | -73.416667 | A | ws | 99 | NYBG | 2394336 | Vasquez | 6308 |
| 124 | *P. subserratum* | no | -3.916667 | -73.583333 | A | ws | 117 | NYBG | 2394339 | Pipoly | 12597 |
| 135 | *P. subserratum* | no | -3.8 | -73.35 | A | ws | 89 | MO | 4395555 | Grandez | 1515 |
| 138 | *P. subserratum* | yes | -3.83661 | -73.36975 | A | ws | 97 | MO | 4891565 | Grandez | 4582 |
| 139 | *P. subserratum* | yes | -3.83661 | -73.36975 | A | ws | 97 | MO | 4891567 | Grandez | 4666 |
| 143 | *P. subserratum* | no | -3.8 | -73.416667 | A | ws | 99 | NYBG | 1241760 | Vasquez | 7594 |
| 90 | *P. subserratum* | no | -0.090278 | -76.213889 | C | non-ws | 278 | NYBG | 2408688 | Jaramillo | 8486 |
| 95 | *P. subserratum* | no | -0.483333 | -75.533333 | C | non-ws | 205 | MO | 4661119 | Palacios | 7572 |
| 106 | *P. subserratum* | no | -0.1 | -76.166667 | C | non-ws | 236 | MO | 5986407 | Brandbyg | 33764 |
| 110 | *P. subserratum* | no | -2.48 | -73.61 | C | non-ws | 129 | MO | 3280977 | Croat | 20516 |
| 125 | *P. subserratum* | no | -3.333333 | -75.916667 | C | non-ws | 195 | NYBG | 2219383 | Melo | 620A |
| 141 | *P. subserratum* | no | -3.25 | -72.9 | C | non-ws | 119 | MO | 6079301 | Grandez | 3599 |
| 147 | *P. subserratum* | no | -3.783333 | -70.283333 | C | non-ws | 105 | MO | 3878917 | Rudas | 1752 |
| 199 | *P. subserratum* | no | -6.93547 | -75.97514 | D | non-ws | 1166 | UC | Paul Fine: Personal Collection | Shavera | 311YSS |
| 200 | *P. subserratum* | no | -6.93547 | -75.97514 | D | non-ws | 1166 | UC | Paul Fine: Personal Collection | Shavera | 685YSS |
| 201 | *P. subserratum* | no | -6.93547 | -75.97514 | D | non-ws | 1166 | UC | Paul Fine: Personal Collection | Shavera | 222YSS |
| 202 | *P. subserratum* | no | -7.06831 | -76.00936 | D | non-ws | 967 | UC | Paul Fine: Personal Collection | Shavera | 667YSS |
| 203 | *P. subserratum* | no | -7.0655 | -75.99419 | D | non-ws | 691 | UC | Paul Fine: Personal Collection | Shavera | 472YSS |
| 155 | *P. subserratum* | yes | 2.72284 | -67.55123 | F | ws | 111 | US | 1833563 | Williams | 14413 |
| 158 | *P. subserratum* | no | 1.933333 | -67.05 | F | ws | 100 | MO | 2992862 | Clark | 6672 |
| *176* | *P. ferrugineum* | no | -8.000000 | -63.000000 | Out-group | NA | NA | NYBG | 1275047 | Teixeira | 140 |
| 187 | *P. ferrugineum* | no | -7.616667 | -72.916667 | Out-group | NA | NA | NYBG | 865157 | Ferreira et al. | 10968 |

| | | | | | | | | | | | |
|---|---|---|---|---|---|---|---|---|---|---|---|
| 179 | *P. ferrugineum* | no | 1.916667 | -67.033333 | Out-group | NA | NA | NYBG | 1275030 | Aymard | 12646 |
| 180 | *P. ferrugineum* | no | 3.834889 | -77.253056 | Out-group | NA | NA | NYBG | 1413893 | Daly | 13836 |
| 178 | *P. ferrugineum* | no | 3.836139 | -77.252222 | Out-group | NA | NA | NYBG | 1413899 | Daly | 13839 |
| 183 | *P. ferrugineum* | no | 3.983333 | -77.033333 | Out-group | NA | NA | NYBG | 1275019 | Gentry | 56707 |
| 181 | *P. ferrugineum* | no | 3.836111 | -77.252222 | Out-group | NA | NA | NYBG | 1413894 | Daly | 13838 |
| 208 | *P. amazonicum* | no | | | Out-group | NA | NA | UC | Paul Fine: Personal Collection | Fine | |

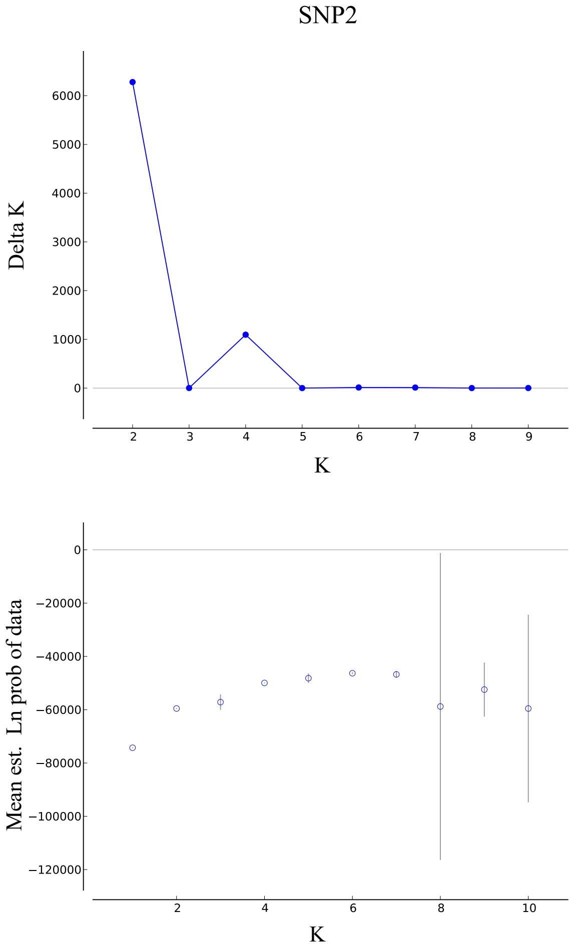


**Figure S1**. Delta K and mean log probability of the data for dataset SNP2, K= 1-10.

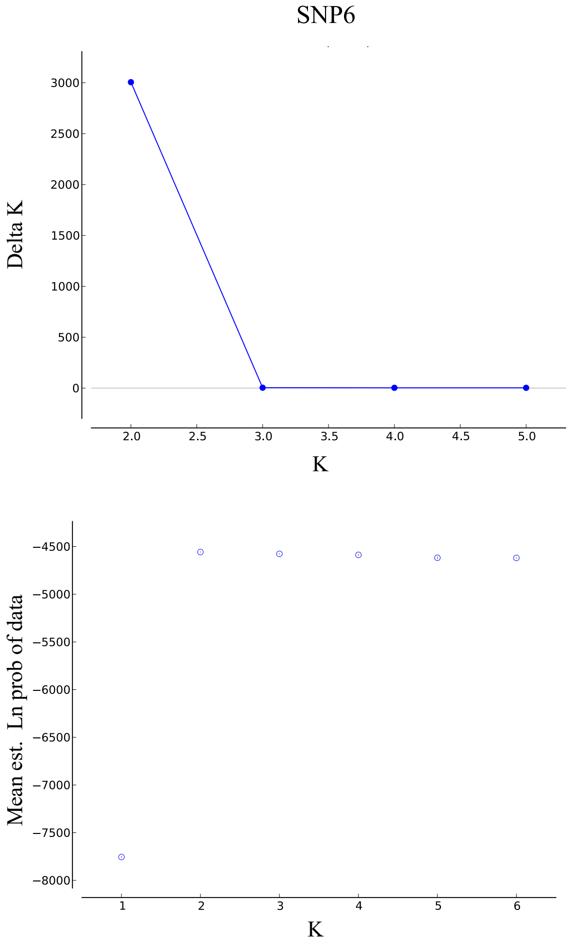


**Figure S2**. Delta K and mean log probability of the data for dataset SNP6, K= 1-6.

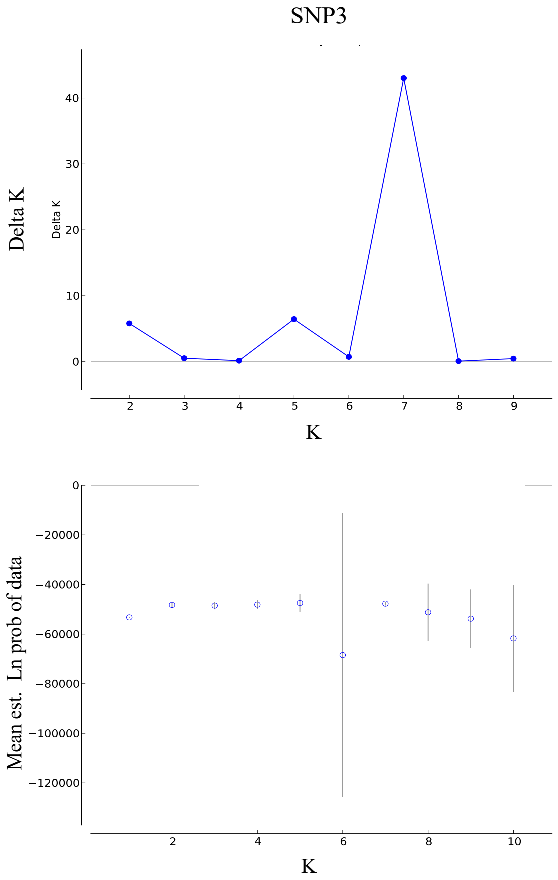


**Figure S3**. Delta K and mean log probability of the data for dataset SNP3, K= 1-10.

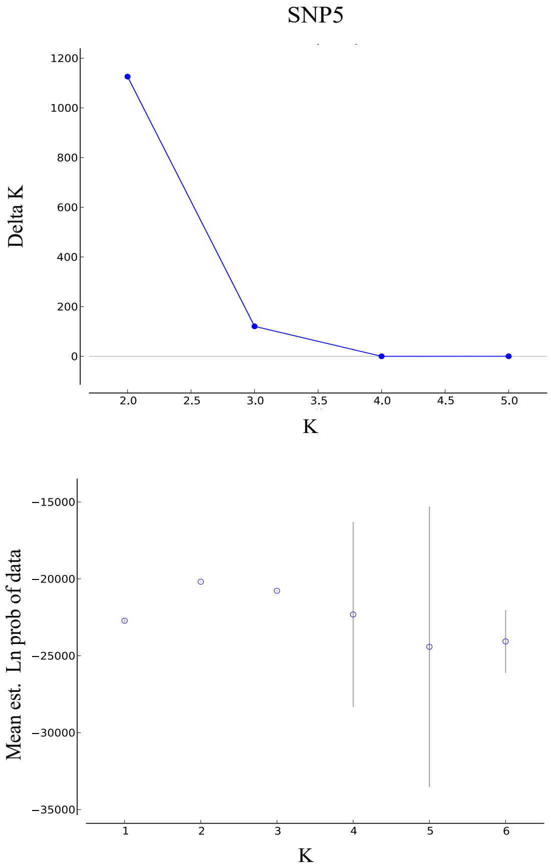


**Figure S4**. Delta K and mean log probability of the data for dataset SNP5, K= 1-6.

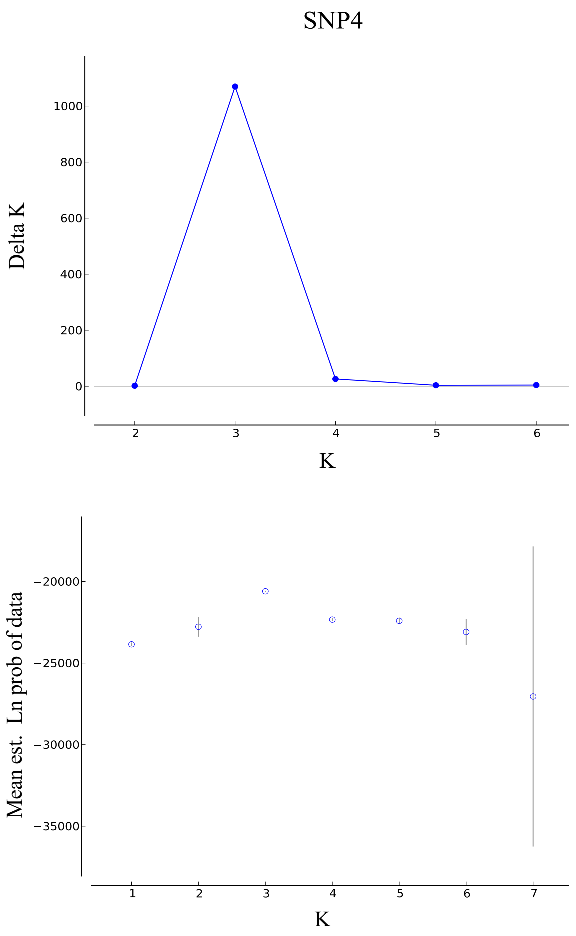


**Figure S5**. Delta K and mean log probability of the data for dataset SNP4, K= 1-7.

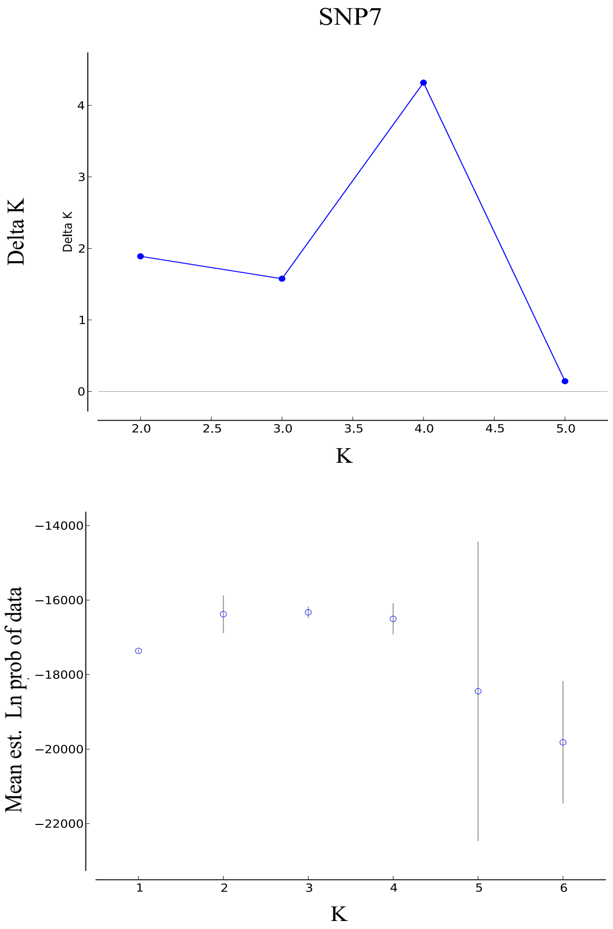


**Figure S6**. Delta K and mean log probability of the data for dataset SNP7, K= 1-6.

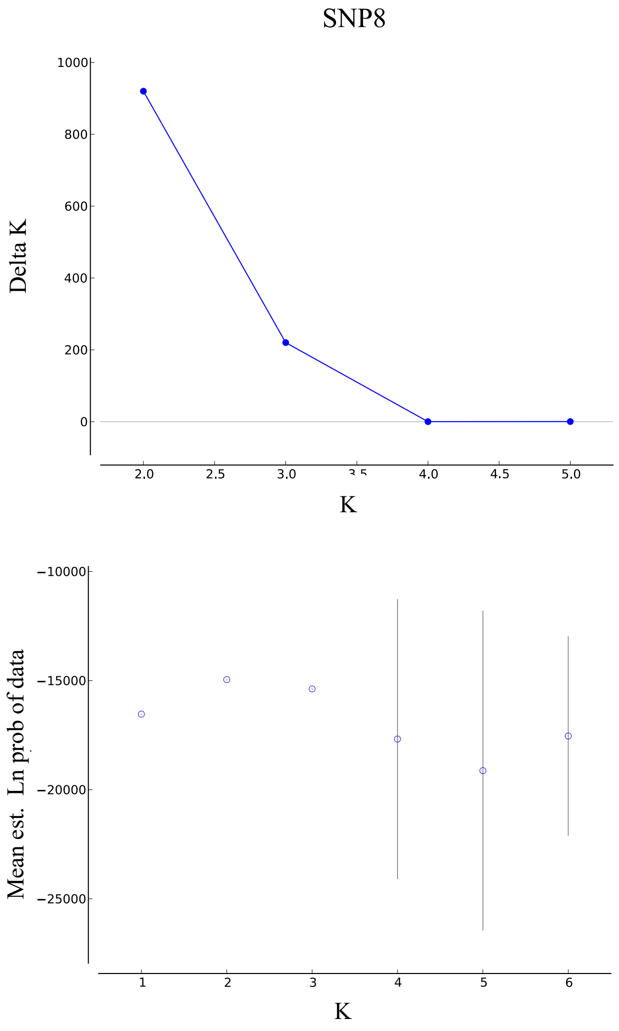


**Figure S7**. Delta K and mean log probability of the data for dataset SNP8, K= 1-6.

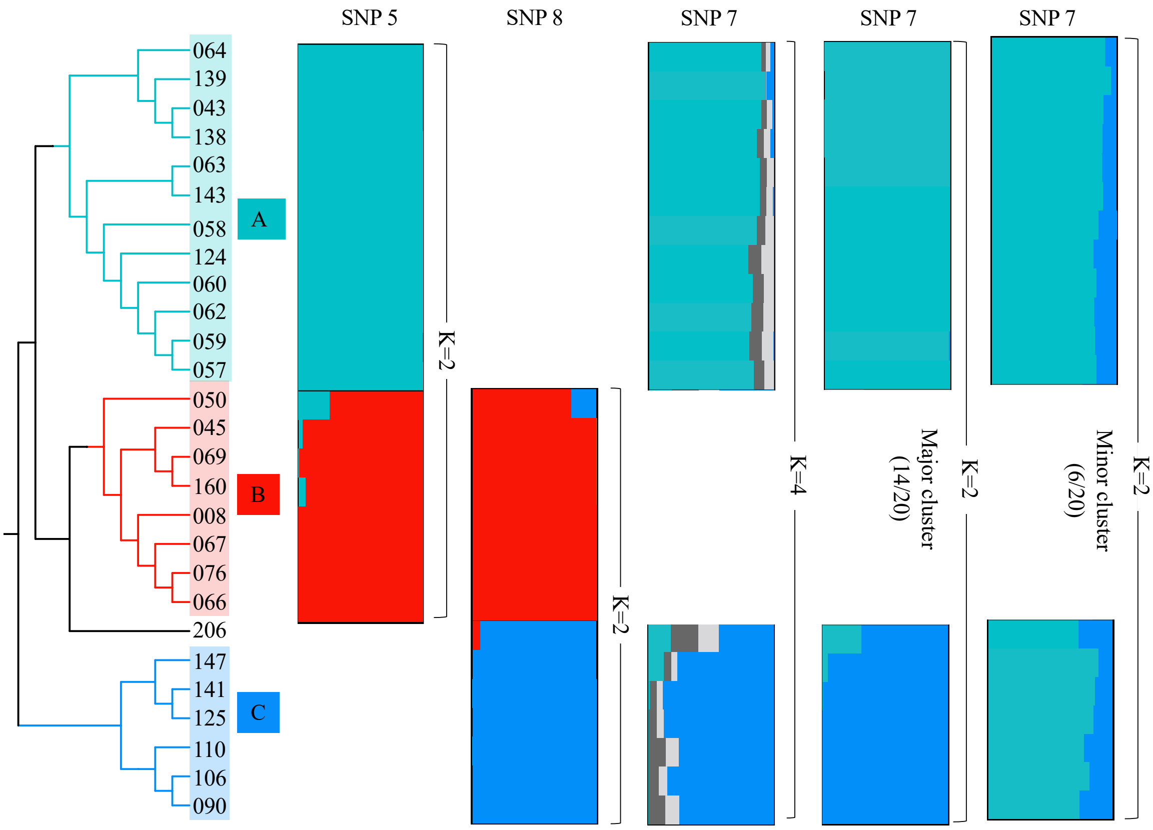


**Figure S8.** Best supported models from three Structure analyses using datasets SNP5, SNP7, and SNP8.

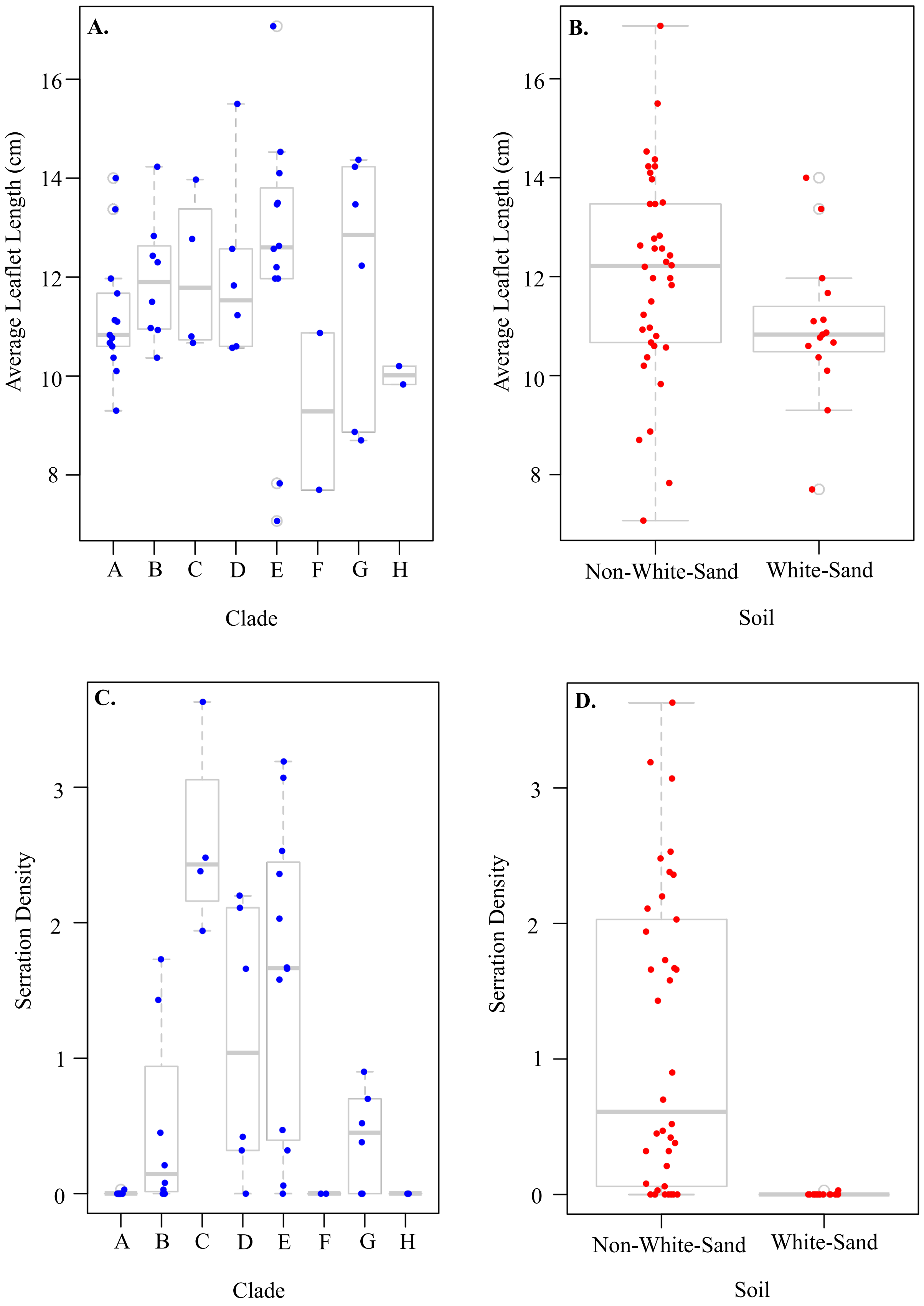
A.
B.
C.
D.
Average Leaflet Length (cm)
16
14
12
10
8
A B C D E F G H
Clade
Non-White-Sand
White-Sand
Soil
Serration Density
3
2
1
0

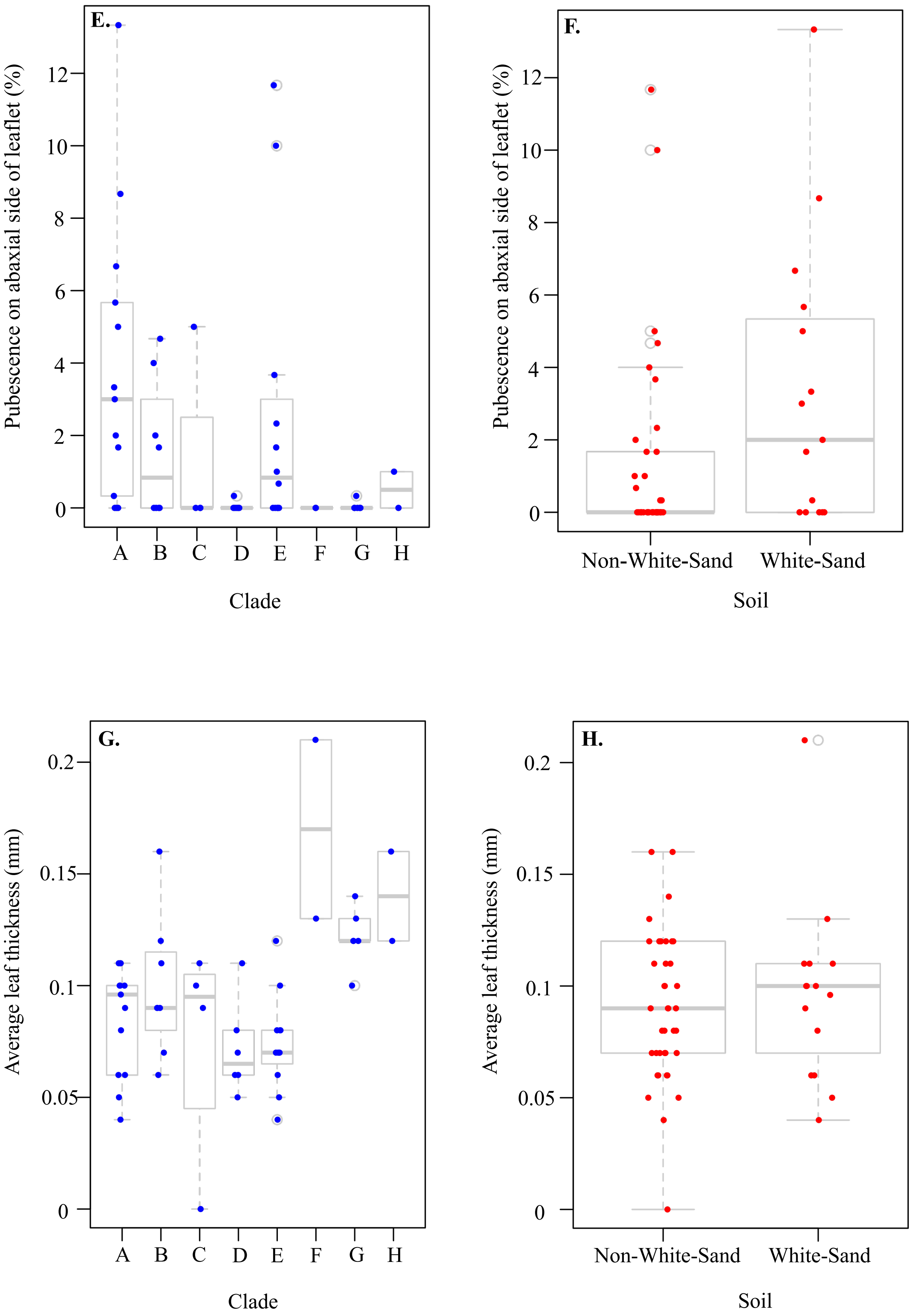
E.
Pubescence on abaxial side of leaflet (%)
0
2
4
6
8
10
12
A
B
C
D
E
F
G
H
Clade
F.
Pubescence on abaxial side of leaflet (%)
0
2
4
6
8
10
12
Non-White-Sand
White-Sand
Soil
G.
Average leaf thickness (mm)
0
0.05
0.1
0.15
0.2
A
B
C
D
E
F
G
H
Clade
H.
Average leaf thickness (mm)
0
0.05
0.1
0.15
0.2
Non-White-Sand
White-Sand
Soil

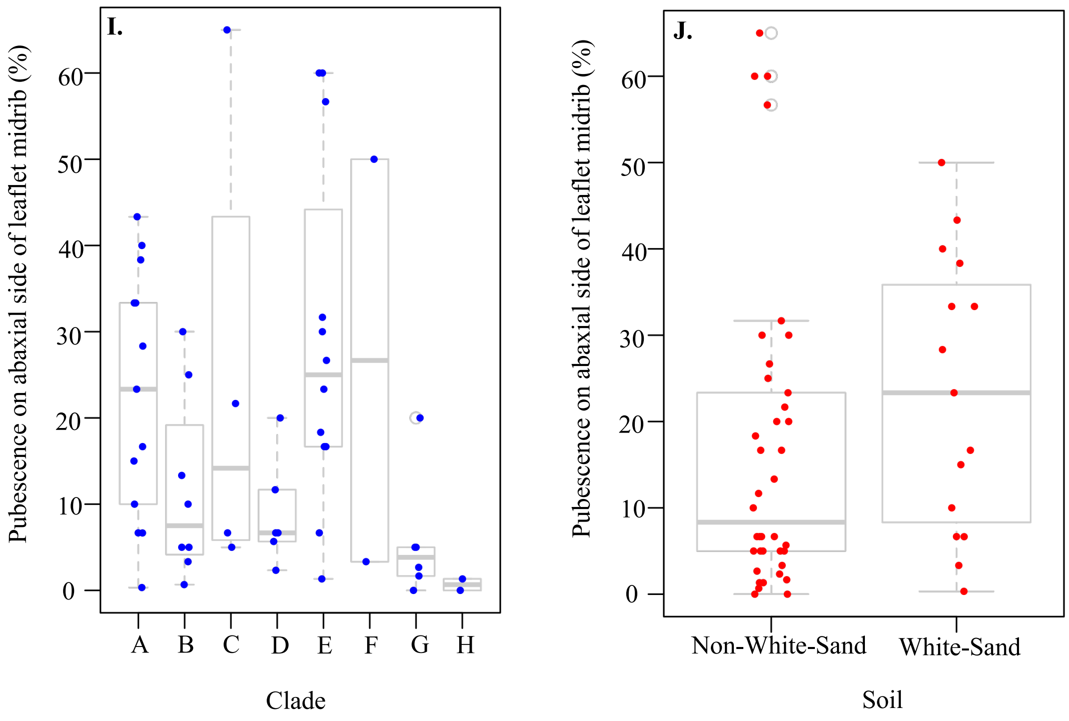


**Figure S9.** Box-whisker plots for each morphological character by clade and soil type. Each point represents an individual. **A**. Average leaf length by clade. **B**. Average leaf length by soil type **C**. Serration density by clade**. D.** Serration density by soil type **E.** Pubescence on the abaxial side of the leaflet blade (%) by clade. **F**. Pubescence on the abaxial side of the leaflet blade by soil type. **G.** Average leaflet thickness by clade. **H.** Average leaflet thickness by soil. **I.** Pubescence on abaxial side of the leaflet midrib (%) by clade. **J.** Pubescence on abaxial side of the leaflet midrib (%) by clade.
